# Single-cell allele-specific expression analysis reveals dynamic and cell-type-specific regulatory effects

**DOI:** 10.1101/2022.10.06.511215

**Authors:** Guanghao Qi, Benjamin J. Strober, Joshua M. Popp, Hongkai Ji, Alexis Battle

**Author notes:** Correspondence to Alexis Battle.

## Abstract

Allele-specific expression, which measures the expression of two alleles of a gene in a diploid individual, is a powerful signal to study cis-regulatory effects. Comparing ASE across conditions, or differential ASE, can reveal context-specific gene regulation. Recently, single-cell RNA sequencing (scRNA-seq) has allowed the measurement of ASE at the resolution of individual cells, but there is a lack of statistical methods to analyze such data. We develop DAESC, a statistical method for differential ASE analysis across any condition of interest using scRNA-seq data from multiple individuals. DAESC includes a baseline model based on beta-binomial regression with random effects accounting for multiple cells from the same individual (DAESC-BB), and an extended mixture model that incorporates implicit haplotype phasing (DAESC-Mix). We demonstrate through simulations that DAESC accurately captures differential ASE effects in a wide range of scenarios. Application to scRNA-seq data from 105 induced pluripotent stem cell lines identifies 657 genes that are dynamically regulated during endoderm differentiation. A second application identifies several genes that are differentially regulated in pancreatic endocrine cells between type 2 diabetes patients and controls. In conclusion, DAESC is a powerful method for single-cell differential ASE analysis and can facilitate the discovery of context-specific regulatory effects.

## Introduction

Allele-specific expression (ASE) measures the expression of one parental allele of a gene relative to the other in a diploid individual. ASE is a powerful tool to study allelic imbalance caused by cis-regulatory genetic variation^1–3^ and epigenetic alterations such as imprinting^4^. In particular, expression quantitative trait loci (eQTL) in or near a gene can cause two alleles to be expressed at different levels^1,2^. Compared to standard eQTL testing, ASE is less susceptible to some confounders including environmental and technical conditions. In addition, comparison of ASE across conditions (differential ASE or D-ASE) can reveal context-specific cis-regulatory effects. Previous ASE studies found that regulatory effects can vary by smoking status, blood pressure medication usage^5^, stages of CD4+ T cell activation^6^, etc.

ASE has been extensively explored using bulk RNA sequencing, but this cannot capture heterogeneity across cell types withing a tissue. Recently, single-cell RNA sequencing (scRNA-seq) has enabled the quantification of ASE at the resolution of individual cells^7–10^ (**Figure 1a**), often across multiple individuals. In this paper, we focus on identifying genes that show differential ASE across conditions. Such methods are only beginning to emerge and are currently applicable to a limited set of scenarios due to assumptions of the models^11,12^. scDALI^11^ uses a beta-binomial mixed-effects model to detect differential allelic imbalance across discrete cell types or continuous cell states. Another method, airpart^12^, partitions the data into groups of genes and cells with similar patterns of allelic imbalance. Airpart also has a function for differential ASE testing based on a hierarchical Bayesian model^12^.

**Figure 1.**
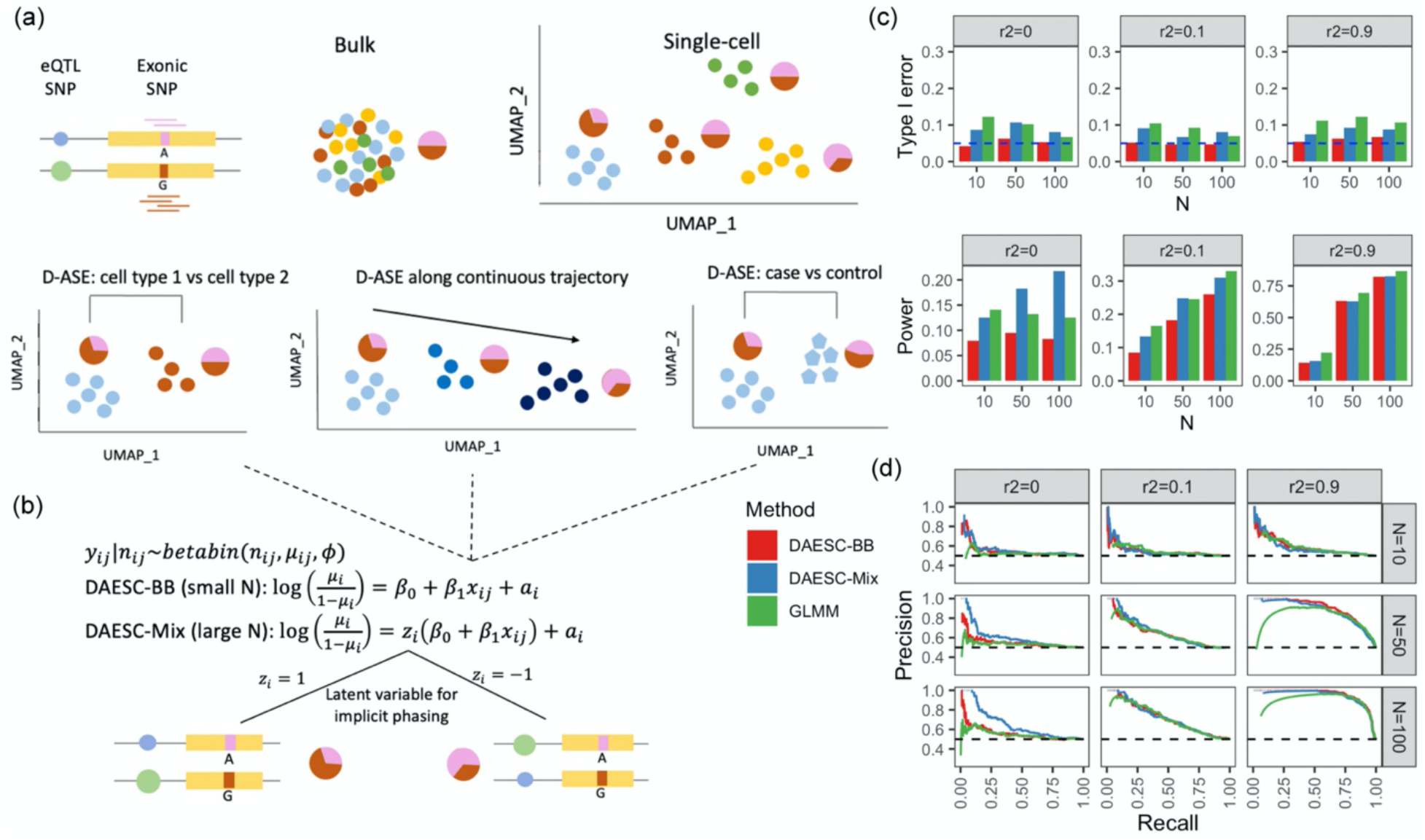
Schematic of DAESC and simulation studies. A) Schematic of allelespecific expression (ASE) measured in bulk tissue and single cells, and three types of differential ASE analysis. Pie charts represent the relative expression of two alleles. b) DAESC models. DAESC accounts for sample repeat structure (multiple cells per sample) using random effects *a_i_* and implicit haplotype phasing using latent variables *z_i_*. c) From simulations, type I error and power under significance threshold p<0.05 and d) precision-recall curves for differential ASE detection along a continuous variable observed in simulations. Allele-specific read counts are simulated from beta-binomial mixture model assuming only one eQTL drives ASE at an exonic SNP. The linkage disequilibrium between the eQTL and the exonic SNP is varied to *r*^2^ = 0,0.1,0.9, and the sample size (number of individuals) is varied to N=10, 50, 100.

However, scDALI or airpart is not optimized for analyzing scRNA-seq data of multiple individuals. One major challenge that is not addressed is how to align read counts consistently across individuals. In the eQTL setting, for example, the eQTL that drives the ASE is not observed. Its expression-increasing allele can be on the haplotype of either the alternative or the reference allele of the exonic SNP where ASE is assessed (eSNP, **Figure 1b**)^5,13,14^. As a result, different individuals may have opposite allelic imbalance actually representing the same regulatory effect. We refer to this phenomenon as “haplotype switching” in the rest of the paper. If not addressed, allelic imbalance will cancel each other across individuals, leading to diminished signal. This issue also exists for ASE caused by epigenetic factors. Previous cross-individual ASE methods for bulk RNA-seq use a majority voting approach, which treats the lower allelic read count as the alternative allele read count^5,14^. This approach, however, is not applicable to single-cell ASE due to low total read count per cell. The scDALI paper avoided this issue with an extra step in the preprocessing, by using phased genotype data and pre-identified eQTLs to align read counts^11^. This approach is not applicable to general differential ASE settings where genotypes are not necessarily measured, or if no significant eQTL is already identified for the gene. A second challenge arising from scRNA-seq data of multiple individuals is the sample repeat structure caused by having multiple cells per individual. This can cause false positives if all cells are treated as independent^11^. scDALI and airpart can account for this structure by adjusting donor IDs as fixed-effects covariates^11,12^. However, this approach is not applicable to comparing ASE between groups of individuals, e.g., disease cases vs controls, since donor IDs as fixed effects can cause collinearity with the binary variable of disease status.

We develop Differential Allelic Expression using Single-Cell data (DAESC), a statistical framework for identifying genes with differential ASE using scRNA-seq data of multiple individuals. DAESC accounts for haplotype switching using latent variables and sample repeat structure of single-cell data using random effects. Simulations studies show the method has robust type I error and high power for differential ASE testing. Applied to single-cell ASE data of 105 individuals^10^, DAESC identifies hundreds of genes with dynamic ASE during endoderm differentiation. A second application to a smaller dataset^8^ identifies 3 genes with differential ASE in pancreatic endocrine cells between type 2 diabetes (T2D) patients and controls.

## Results

### Overview of DAESC

DAESC is based on beta-binomial regression model and can be used for differential ASE against any independent variable *x_ij_*, such as cell types, continuous developmental trajectories, genotype (eQTLs), or disease status (**Figure 1a**). DAESC is comprised of two components (DAESC-BB and DAESC-Mix) to be used under different scenarios (**Figure 1b**). The baseline model DAESC-BB is a beta-binomial model with individual-specific random effects (*a_i_*) that account for the sample repeat structure (**Methods**), arising from multiple cells measured per individual. DAESC-BB can be used generally for differential ASE regardless of sample size. When sample size is reasonably large (e.g. N≥20, we introduce a full model DAESC-Mix that accounts for both sample repeat structure and implicit haplotype phasing (**Methods**). For example, when ASE measured at a heterozygous exonic SNP (eSNP) is driven by an eQTL, the expression-increasing allele of the eQTL could be on the haplotype of the alternative allele of the eSNP (*z_i_* = 1), or the reference allele of the eSNP (*z_i_* = −1). We account for this possibility using latent variables *z_i_*, which lead to a mixture model (**Figure 1b**). Though it is possible that the true model may have more mixture components especially when the gene has multiple eQTLs, we use the two-component mixture model to prevent against overfitting and increase computational speed. For both DAESC-BB and DAESC-Mix, parameter estimation is conducted using variational EM algorithm (see **Methods** and **Supplementary Notes** for details). Hypothesis testing for differential ASE (*H*_0_: *β*_1_ = 0) is conducted using likelihood ratio test.

### Simulation studies

We first conduct simulations from beta-binomial mixture model assuming only one eQTL drives the ASE at the eSNP. In the first scenario where we test differential ASE along a continuous variable representing cell state (e.g., differentiation stage), we observe that DAESC-BB has well-controlled type I error across scenarios (**Figure 1c**). DAESC-Mix has slight type I error inflation (averaged 8.5% across scenarios) but less than a standard GLMM (averaged 10% across scenarios). When there is no LD between the eQTL and eSNP (r^2^=0), we observe a substantial power gain by using DAESC-Mix compared to DAESC-BB and GLMM. The gain is more pronounced when the sample size is large (N=50 or 100). This is likely due to the ability of DAESC-Mix to conduct implicit haplotype phasing. When r^2^=0.1, DAESC-Mix has similar power to GLMM, and both are slightly more powerful than DAESC-BB. When the LD between the eQTL and eSNP is strong (r^2^=0.9), we observe only minimal power difference across the three methods. Results from the GTEx Consortium^15^ show LD r^2^<0.1 for most eQTL-eSNP pairs (**Supplementary Figure 1**), indicating that for most genes DAESC-Mix is likely to lead to improved power. The precision-recall curves show that DAESC-Mix dominates the other two methods when r^2^=0 and *N* ≥ 50 with varying significance thresholds (**Figure 1d**). In addition, the curves for GLMM tend to dip near low recall value, i.e., when the significant threshold is stringent. This indicates potential issues with p-value calibration for GLMM.

For differential ASE with respect to binary case-control disease status, we observe mostly similar patterns as those in the previous simulation with continuous cell state (**Supplementary Figure 2**). A notable distinction is that all methods have more inflated type I error (~10%) when *N* ≤ 10, and GLMM has higher type I error inflation across scenarios. The pseudobulk-based method, EAGLE-PB, has similar performance with DAESC-BB except when r^2^=0.9, where DAESC-BB appears slightly more powerful (**Supplementary Figure 2**). EAGLE-PB assumes independent samples and is not applicable to the continuous-cell-state simulations shown in Figure 1.

Since eQTL studies have found that allelic heterogeneity is widespread^16–19^, we also investigate the performance of the methods when there are multiple eQTLs driving the ASE. Due to the large number of scenarios for the LD across multiple eQTLs and the eSNP, here we only investigate the scenario where no LD exists among the eQTLs or with the eSNP. Similar to the previous scenario, DAESC-BB controls type I error under varying number of eQTLs; DAESC-Mix, though having slightly inflated type I error in some settings, is less inflated than GLMM (**Figure 2a**). This shows that although multiple eQTLs introduces extra mixture components into the true model (**Methods**), it has minimal impact on the type I error control. In addition, we observe a substantial power gain by DAESC-Mix compared to DAESC-BB or GLMM (**Figure 2a**), which is more pronounced than when only one eQTL drives ASE (**Figure 1**). This gain exists not only under large sample size, but also under small sample size (N=10) despite a smaller margin. In addition, power increases steadily for DAESC-Mix with increasing number of eQTLs, showing larger advantage over DAESC-BB and GLMM under allelic heterogeneity (**Figure 2a**). Precision-recall curves show that DAESC-Mix consistently outperforms the other two methods across different significance thresholds, with DAESC-BB ranking second (**Figure 2b**).

**Figure 2.**
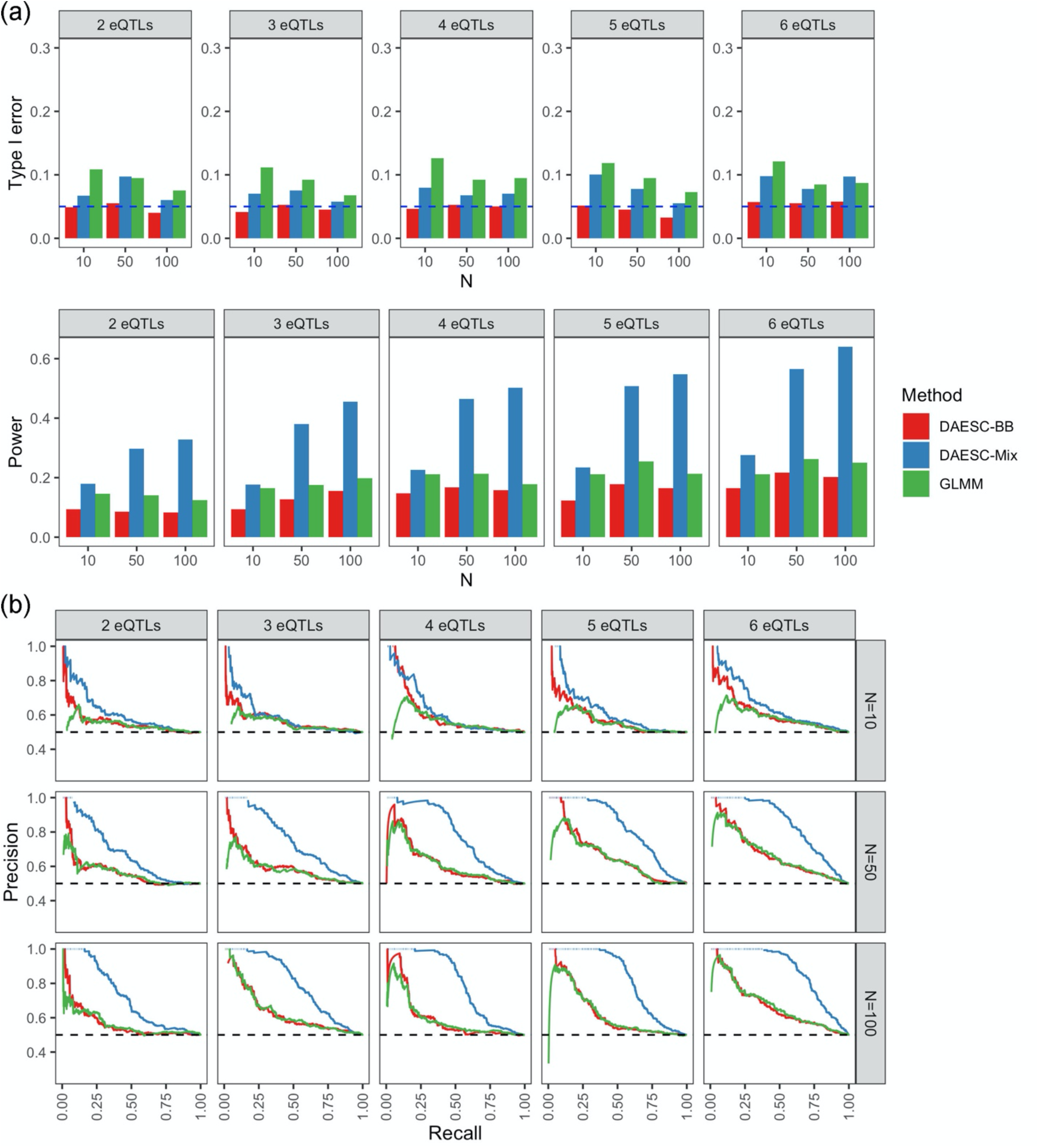
Simulation studies with multiple eQTL SNPs per gene. A) Type I error and power and b) precision-recall curves for differential ASE detection along a continuous variable observed in simulations. Allele-specific read counts are simulated from betabinomial mixture model assuming multiple eQTLs drives ASE of an exonic SNP. We assume no linkage disequilibrium among the eQTLs and exonic SNP. The sample size (number of individuals) is varied N=10, 50, 100.

When testing differential ASE for binary case-control disease status, DAESC-Mix remains most powerful when there are multiple eQTLs per eSNP (**Supplementary Figure 3**). In fact, DAESC-BB, GLMM and EAGLE-PB, which do not conduct implicit phasing, do not appear to have any power to detect differential ASE. In contrast to D-ASE along continuous cell state (**Figure 2**), the power of DAESC-Mix changes minimally the number of eQTLs (**Supplementary Figure 3**). This indicates that cell-level variability, which is a special feature of single-cell ASE, could be important for implicit phasing.

### Dynamic ASE during endoderm differentiation

We apply DAESC-BB, DAESC-Mix and GLMM to single-cell ASE data for 30,474 cells from 105 individuals collected by Cuomo et al^10^. In their experiment, induced pluripotent stem cells (iPSCs) underwent differentiation for three days into mesendoderm and definitive endoderm cells (**Figure 3a**). To study dynamic regulatory effects along the differentiation trajectory, we conduct differential ASE analysis along pseudotime (*x_ij_*), which was estimated and provided by the original study (**Figure 3b**). DAESC-BB identifies 324 dynamic ASE (D-ASE) genes that vary along pseudotime, and DAESC-Mix identifies 657 D-ASE genes (FDR<0.05, **Figure 3c** and **Supplementary Table 1**). Nearly all genes identified by DAESC-BB are also identified by DAESC-Mix (**Figure 3d**). Since dynamic ASE can be driven by dynamic cis-regulatory effects, we use the overlap between our D-ASE genes and dynamic eGenes reported by Cuomo et al^10^ as a validation criterion. Among the genes identified by DAESC-BB, 35.5% were reported by Cuomo et al, while 27.5% identified by DAESC-Mix were reported (**Figure 3c**). GLMM identifies a large number D-ASE genes (1,995, FDR<0.05), but have low validation rate (13.4%), indicating potential type I error inflation. Comparing the same number of top genes (smallest p-values) selected by three methods, DAESC-Mix shows higher validation rate than DAESC-BB or GLMM across thresholds (**Figure 3e**). In addition, dynamic ASE genes discovered using DAESC-Mix display total expression dynamics similar to those of previously discovered dynamic eGenes (**Supplementary Figure 4**). This shows that DAESC-Mix offers an increase in power without biasing discovery toward particular trends in expression or technical factors influencing total expression levels.

**Figure 3.**
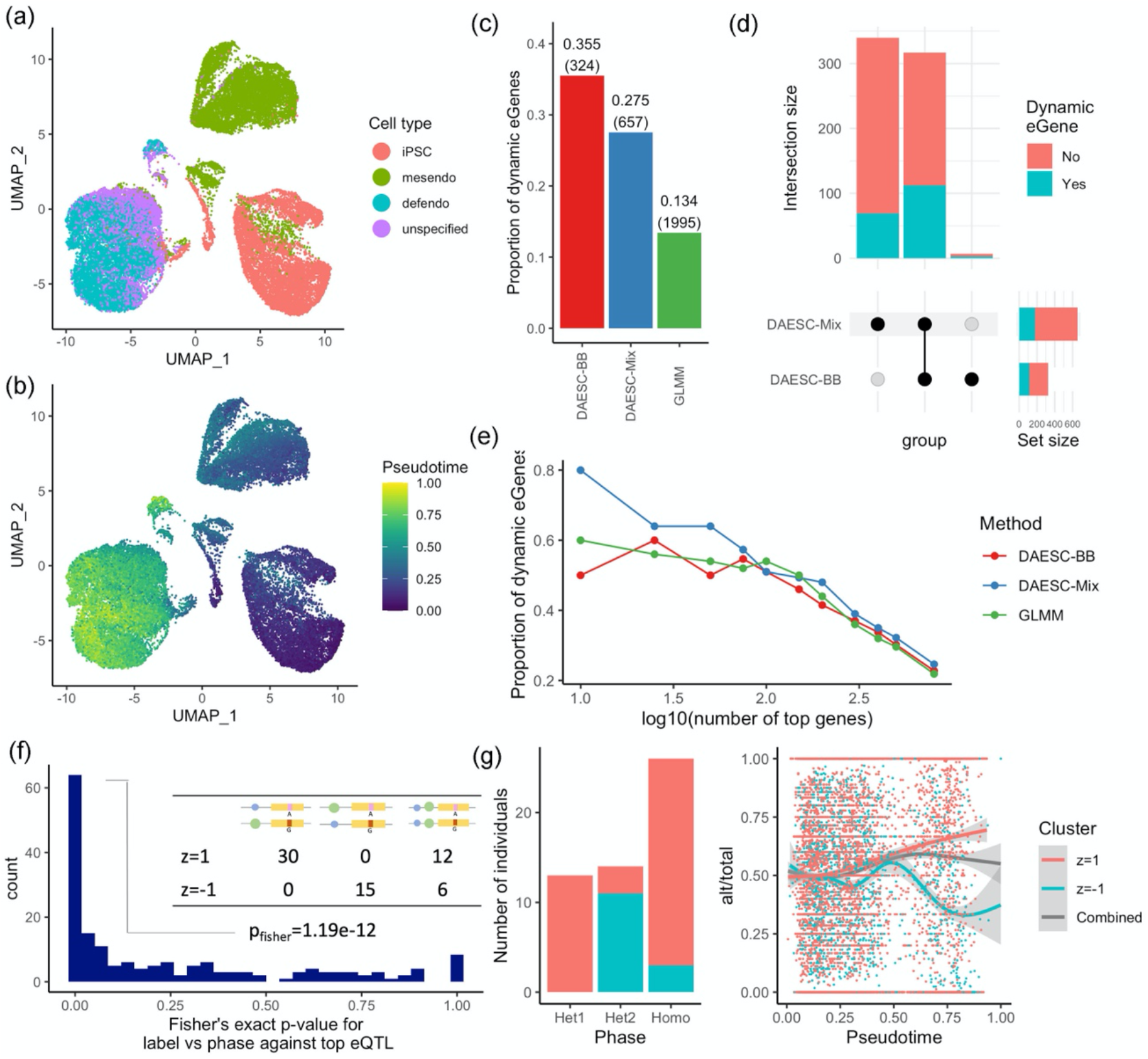
Dynamic ASE during endoderm differentiation. UMAP plot colored by a) cell type and b) pseudotime. Cell types include induced pluripotent stem cells (iPSCs), mesendoderm cells (mesendo) and definitive endoderm cells (defendo). c) Proportion of dynamic ASE (D-ASE) genes identified by three methods that were also dynamic eGenes reported by Cuomo et al. Number of D-ASE genes identified by each method are annotated in the parentheses. d) Number of D-ASE genes identified by DAESC-Mix but not DAESC-BB (first bar), both methods (second bar), and DAESC-BB but not DAESC-Mix. e) Proportion of dynamic eGenes reported by Cuomo et al among varying number of top D-ASE genes identified by three methods. f) Fisher’s exact test p-values testing whether DAESC-Mix cluster labels capture haplotype information between the top exonic SNP and top eQTL reported by Cuomo et al. Schematics of three haplotype combinations are used as column names of the example 2 × 3 table (from left to right: het1, het2, homo). Green and blue circles are the reference (ref) and alternative (alt) alleles of the eQTL, respectively; red and pink rectangles are the alt and ref for the exonic SNP, respectively. g) An example (*NMU* gene) of mixture clusters capturing haplotype information. Alt: alternative allele read count; total: total allele-specific read count.

We further use the phased genotype data to validate the ability of DAESC-Mix to conduct implicit haplotype phasing. We conduct the validation on the genes that show suggestive evidence of D-ASE by DAESC-Mix (p<0.05) and have at least one eQTL reported by Cuomo et al^10^. We further restrict to 179 genes that have significant likelihood ratio test comparing DAESC-Mix to DAESC-BB (p<0.05). This restriction in effect selects genes for which DAESC-Mix reports two haplotype combinations (*z_i_* = 1 and *z_i_* = −1). Fisher’s exact test shows that for 77 (43%) genes, the mixture labels given by DAESC-Mix successfully capture observed haplotype combinations between the gene and the top eQTL (p<0.05, **Figure 3f**). For 39 (22%) genes, mixture labels are not associated with haplotype combinations (p>0.5). This could be due to imperfect eQTL calling by the original study, or limitations of our method. An example is *NMU*, for which DAESC-Mix reports highly significant dynamic ASE (*p* = 1.93 × 10^−59^) and captures the haplotype combinations (*p_fisher_* = 1.51 × 10^−6^). We observe that allelic fractions move in opposite directions along pseudotime for two clusters of individuals, and combining two groups would severely diminish the allelic imbalance (**Figure 3g**).

Due to its high power and validation rate, and ability to capture haplotype combinations, we choose DAESC-Mix as the main method of discovery.

### Patterns and mechanisms of dynamic ASE

We hypothesize that dynamic ASE during differentiation could be linked to dynamic changes of chromatin state. To test this hypothesis, we use the chromatin states learned by ChromHMM^20^ on the Roadmap Epigenomics data^21^ (see **Methods** for details). We recode the chromatin states to 0 (inactive) and 1 (active) based on the criteria described in **Methods**. For each gene, we compute the absolute value of change in chromatin state (0 – inactive, 1 – active) at the transcription start site between two endpoints of differentiation: iPSC and definitive endoderm. The D-ASE genes identified by DAESC-Mix show an average chromatin state change of 0.132, while the non-D-ASE genes show an average change of 0.075 (**Figure 4a**). This difference is highly significant even after adjusting for the read depth of the genes (*p* = 3.19 × 10^−9^). The D-ASE genes identified by DAESC-BB and GLMM also show larger change in chromatin state compared to non-D-ASE genes, but the difference is smaller (**Figure 4a**). The patterns persist if we compare the same number of top genes, instead of the statistically significant ones, identified by each method (**Supplementary Figure 5**). In addition, we observe significant correlations between the D-ASE effect size (log-OR when pseudotime changes from 0 to 1) and the magnitude of change in chromatin state, with DAESC-Mix showing the strongest correlation (**Figure 4b**). Gene-set enrichment analysis found 121 Gene Ontology (GO) biological process gene sets enriched in D-ASE genes identified by DAESC-Mix, including those for the regulation of mesoderm development and cell development (**Supplementary Table 2**).

**Figure 4.**
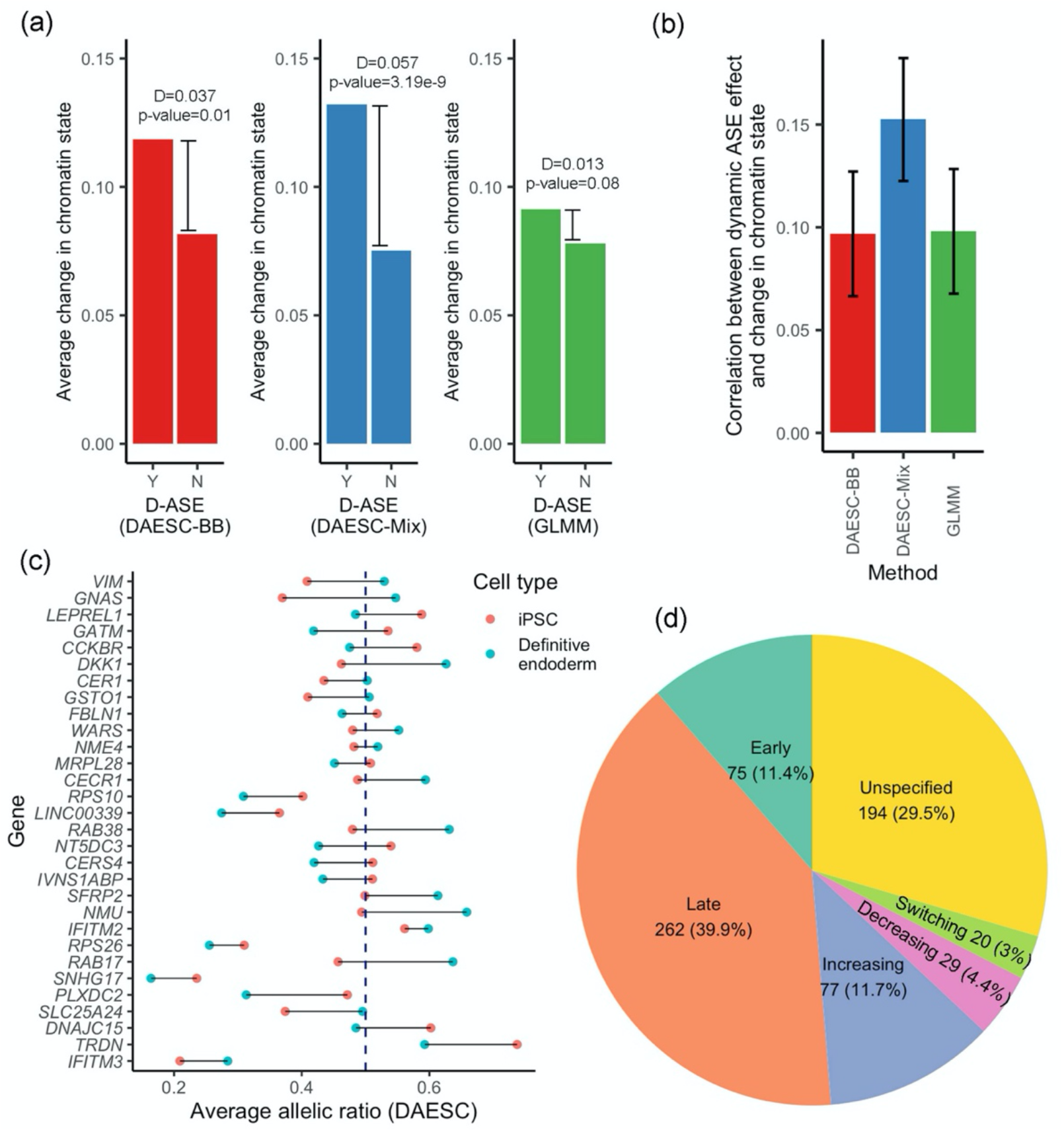
Patterns and mechanisms of dynamic ASE genes during endoderm differentiation. a) Average change of chromatin state at transcription start site from iPSC to definitive endoderm cells for D-ASE genes (Y) and non-D-ASE genes (Y). At FDR<0.05, the number of D-ASE genes (Y) is 324 for DAESC-BB, 657 for DAESC-Mix, and 1,995 for GLMM. For each method, the genes that do not reach FDR<0.05 are considered non-D-ASE genes (see Methods for details). Chromatin states are from ChromHMM analysis of the Roadmap Epigenomics data and recoded to 0 (inactive) or 1 (active). D is the difference between D-ASE and non-D-ASE genes, and p-values are calculated using linear regression: chromatin state change ~ Y/N + total read depth of the gene. b) Correlation between D-ASE effect size (*β*_1_) and change in chromatin state. Error bars represent 95% confidence intervals. c) Top 30 genes identified by DAESC-Mix (smallest p-values) and average allelic ratio of iPSCs vs definitive endoderm cells estimated by DAESC-Mix, computed as 1/(1 + exp(−(*β*_0_ + *β*_1_*t*))) where *t* is the average pseudotime of the cell type. d) Types of D-ASE genes and their proportions. See Methods for details.

To further study the pattern of dynamic change in ASE, we compute the average allelic fraction for iPSCs and definitive endoderms using DAESC-Mix estimates (**Methods**). We found different genes show allelic imbalance at different stages of differentiation (**Figure 4c**). For example, genes *SFRP2* and *NMU* have minimal allelic imbalance at the iPSC stage but substantial imbalance at the definitive endoderm stage. On the contrary, genes *VIM* and *LEPREL1* only shows allelic imbalance in iPSCs but not definitive endoderms. For genes *IFITM3, SNHG17* and *TRDN* the allelic imbalance appears at both stages of differentiation but with a different magnitude. Lastly, for genes *RAB17* and *GATM* the allelic fraction switches directions across stages, i.e., the highly expressed allele for iPSCs becomes the less expressed allele for definitive endoderms. Based on these observations, we classify the 657 D-ASE genes identified by DAESC-Mix into 6 categories based which differentiation stage shows allelic imbalance (**Figure 4d**). More than half of the genes show stronger allelic imbalance in definitive endoderms than iPSCs (51.6% late and increasing, **Figure 4d**), only 15.8% shows stronger imbalance in iPSCs (early and decreasing).

### Type 2 diabetes and differential ASE in pancreatic islet cells

We obtain the scRNA-seq data from pancreatic islet samples of 4 type 2 diabetes (T2D) patients and 6 controls^8^. After preprocessing (**Methods**), we obtain single-cell ASE data for 2,209 cells of >10 cell types (**Figure 5a-b**). To identify genes potentially dysregulated in T2D patients, we conduct differential ASE analysis between cases and controls for four major endocrine cell types: alpha, beta, delta, and gamma cells. Due to the small sample size, we use DAESC-BB as the method for discovery. We found three genes that show differential ASE between cases and controls (FDR<0.05, **Figure 5c**). Among them, the D-ASE of *ARPC1B, SLC37A4* is only found in alpha cells, and the D-ASE of *REEP5* is found in both alpha and beta cells. *SLC37A4* and *REEP5* show stronger allelic imbalance in T2D patients than controls (**Figure 5c**), indicating regulatory effects that are only present in T2D patients. *ARPC1B*, however, shows stronger allelic imbalance in healthy controls (**Figure 5c**), indicating regulatory effects potentially disabled in T2D patients.

**Figure 5.**
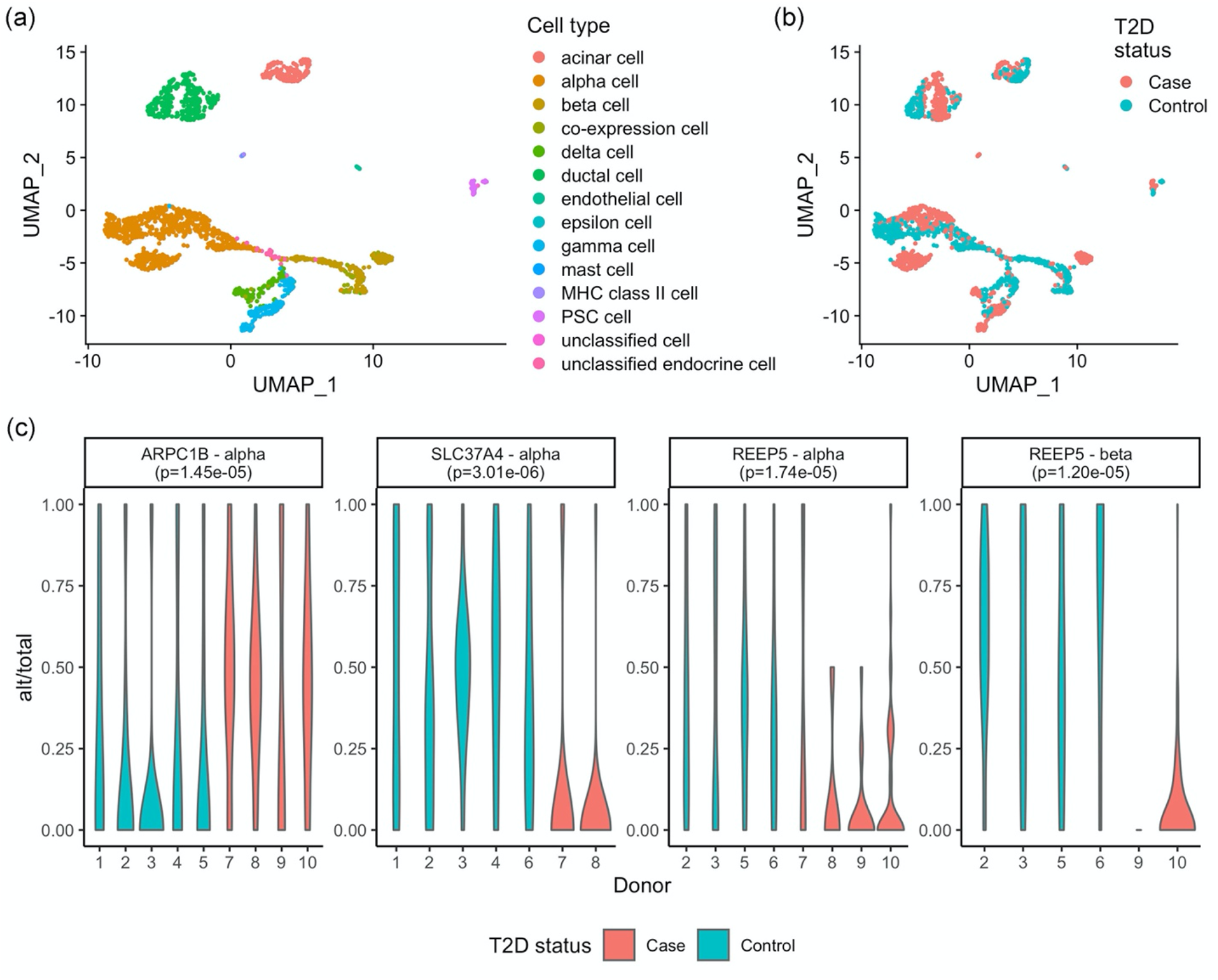
Differential ASE between type 2 diabetes patients and controls in pancreatic endocrine cells. UMAP colored by a) cell type b) disease status. c) Three D-ASE genes in two cell types identified by DAESC-BB and distribution of allelic fraction in each individual donor. Alt: alternative allele read count; total: total allele-specific read count.

Previous studies indicated potential link between *SLC37A4* and T2D. *SLC37A4* encodes glucose 6-phosphate translocase, which transports glucose 6-phosphate from the cytoplasm to the endoplasmic reticulum^22,23^. SNP rs7127212, which is 51.6kb from the TSS of *SLC37A4*, was reported to be associated with the risk of T2D by a previous study^24^. We did not find strong functional connection with T2D for *ARPC1B* and *REEP5* in the existing literature.

## Discussion

Differential allele-specific expression is a powerful tool to study context-specific cis-regulatory effects. Single-cell RNA-seq has allowed the study of ASE in heterogeneous cell types within a tissue. However, there is a lack of statistical tools for single-cell differential ASE analysis. In this paper, we describe DAESC, a generic statistical framework for differential ASE detection using scRNA-seq data from multiple individuals. The method captures sample repeat structure of multiple cells per individual using random effects, and DAESC-Mix further refines differential ASE analysis by incorporating implicit haplotype phasing. Simulation studies show the method has well controlled type I error and high power under a wide range of scenarios. Application to single-cell ASE data from an endoderm differentiation experiment identifies hundreds of genes that are dynamically regulated during differentiation. A second application to single-cell data from pancreatic islets identifies 3 genes with differential ASE between T2D patients and controls in alpha and beta cells, despite the small sample size.

Within the DAESC framework, the full model DAESC-Mix is generally more powerful than DAESC-BB. However, we recommend using DAESC-Mix when the number of individuals is reasonably large (e.g., *N* ≥ 20), since the mixture model needs large N to identify different haplotype combinations. Indeed, simulation studies show that power gain is more pronounced under large N (**Figures 1 and 2, Supplementary Figures 2 and 3**). When the sample size is small (e.g. *N* ≤ 10), the overall performance between DAESC-Mix and DAESC-BB is less distinguishable (see precision-recall curves in **Figure 1** and **Supplementary Figure 2**). In that case, we recommend using DAESC-BB which has better type I error control. In our first application, the data from endoderm differentiation are comprised of 105 individuals and hence DAESC-Mix is chosen. In the second application, the pancreatic islet dataset is comprised of only 10 individuals and hence DAESC-BB is chosen.

Note that the two-component mixture model used by DAESC-Mix is a simplifying assumption. When the gene has one eQTL, the true model should have an extra component corresponding to the individuals of whom the eQTL is homozygous. When the gene has multiple eQTLs, the number of mixture components grows exponentially. DAESC-Mix uses a two-component model to prevent against overfitting and increase computational speed. Simulation studies show the performance of DAESC-Mix remains robust when there are multiple eQTLs (**Figure 2** and **Supplementary Figure 3**). This is also due to the limitation of sample size, since the number of individuals in single-cell ASE datasets are not enough to robustly fit a mixture model with many components. More complex mixture models may become viable as more data are collected.

DAESC has important conceptual and technical differences from scDALI and airpart. First, DAESC is designed as a generic tool for differential ASE analysis with respect to any condition, regardless of whether the comparison is between cell-types within an individual or across individuals, and regardless of whether the condition of interest is continuous or discrete. The random effects that account for sample repeat structure is an important component that enables this flexibility. scDALI and airpart focus on differential ASE across cell types, not across samples or individuals. They allow for adjustment of donor IDs as fixed effects but cannot be used for differential ASE across individual-level conditions (e.g. disease status). Due to these distinctions, the GLMM fitted by lme4 is more comparable to DAESC-BB than scDALI and airpart, and hence used as the main reference method for benchmarking. Second, DAESC-Mix further conducts implicit haplotype phasing to recover allelic signals hidden by haplotype switching. Hence DAESC-Mix can be powerful regardless of whether genotypes are available or eQTLs have been identified, which is often not the case for case/control comparisons. In the scDALI paper^11^, the application to scRNA-seq data assigned the alternative haplotype of the gene based on the alternative allele of the eQTL. This approach is only possible if genotype data are available, and there is at least one significant eQTL for the gene. If the gene is regulated by multiple weak eQTLs that do not attain genome-wide significance, scDALI does not have a mechanism to assign alternative haplotypes. However, DAESC-Mix can still be used and may be able to capture the combined effects of multiple eQTLs as shown in the simulations (**Figure 2** and **Supplementary Figure 3**). Previous methods for bulk RNA-seq have used a majority voting approach for pseudo haplotype phasing^5,14,25^. However, this approach is not directly applicable to single-cell ASE due to multiple cells from each individual and low read depth per cell.

Our method does have some limitations to consider. First, we observe modest type I error inflation for DAESC-Mix potentially due to overfitting. However, the inflation seems acceptable given the magnitude of power improvement. If provided with enough computational resources, the users can choose to conduct permutation tests to further correct type I error. Second, DAESC-Mix is most powerful when applied to datasets with a large number of individuals, but such datasets are not widely available. For small datasets we recommend using DAESC-BB, which may be conservative but has well controlled type I error. In the future, DAESC-Mix could be more widely applied with the availability of new technology for large-scale single-cell ASE profiling. Lastly, DAESC is not optimized for integrating information across multiple cell types into a unified test. scDALI and airpart both have methods for this purpose. A future direction is to combine the strengths of DAESC and scDALI or airpart to incorporate sample repeat structure, implicit haplotype phasing and integration of information across cell types.

In conclusion, we develop a statistical method, DAESC, for powerful detection of differential ASE across a wide variety of conditions. DAESC will be one of the first methods for this purpose and has complementary strengths to existing methods.

## Methods

### DAESC model

We describe the DAESC model for differential ASE analysis using scRNA-seq data across multiple individuals. For a heterozygous exonic SNP, let *y_ij_* be the alternative allele read count for individual *i* and cell *j*, and *n_ij_* be the total allele-specific read count. Let *x_ij_* be the independent variable, e.g. cell types, cell differentiation time, or disease status of the individual. Define ***y**_i_* = (*y*_*i*1_, …, *y_iJ_i__*) where *J_i_* is the number of cells from individual *i*. DAESC is comprised of two components: a baseline betabinomial regression model with individual-specific random effects (DAESC-BB), and a full betabinomial mixture model that incorporates implicit phasing (DAESC-Mix).

The DAESC-BB model is formulated as follows

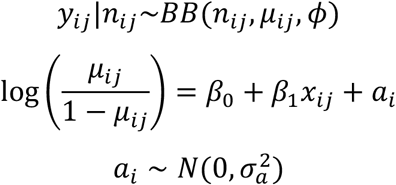

Here *BB*(*n_ij_, μ_ij_, ϕ*) is a beta-binomial distribution with denominator *n_ij_*, mean proportion *μ_ij_* and overdispersion parameter *ϕ*. It is equivalent to 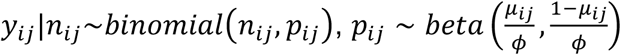 marginalized over *p_ij_*. We model 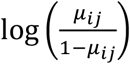 as a linear function of *x_ij_*. The individual-specific random effect *a_i_* accounts for the sample repeat structure introduced by having multiple cells from each individual. This model can be used for any differential ASE analysis but may be conservative in some scenarios due to unknown causal variants and haplotype information. For example, when the exonic SNP is not in strong LD with the causal eQTL, different individuals may exhibit complementary allelic fractions which actually reflect the same regulatory effect. Failing to account for this possibility can lead to diminished ASE signal when aggregated across individuals.

This issue can be addressed using DAESC-Mix when the sample size (number of individuals) is sufficiently large. The model is formulated as follows

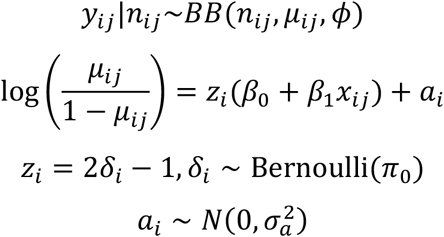

This model is an extension of DAESC-BB with the inclusion of an indicator variable *z_i_*. It models the scenario where ASE is caused by one regulatory SNP (rSNP). When *z_i_* = 1, the alternative allele of the eQTL and the alternative allele of the exonic SNP is on the same haplotype, and the reference alleles of the two SNPs are on the same haplotype. When *z_i_* = −1, the alternative allele of the eQTL and the reference allele of the exonic SNP is on the same haplotype, and vice versa (**Figure 1**). Though it is possible that the eQTL is homozygous for some individuals, we do not model this scenario to protect against overfitting and speed up computation.

Though the models above are described for a heterozygous exonic SNP, it can also be applied to gene-level ASE counts generated by aggregating across multiple exonic SNPs.

### Model inference by variational EM algorithm

The inference is conducted by variational EM algorithm^26^. Here we describe the algorithm for DAESC-Mix. Details of the derivation and the algorithm for DAESC-BB can be found in **Supplementary Notes**. Denote ***β*** = (*β*_0_, *β*_1_)^*T*^. We treat *a_i_* and *δ_i_* as missing data and the complete data likelihood is

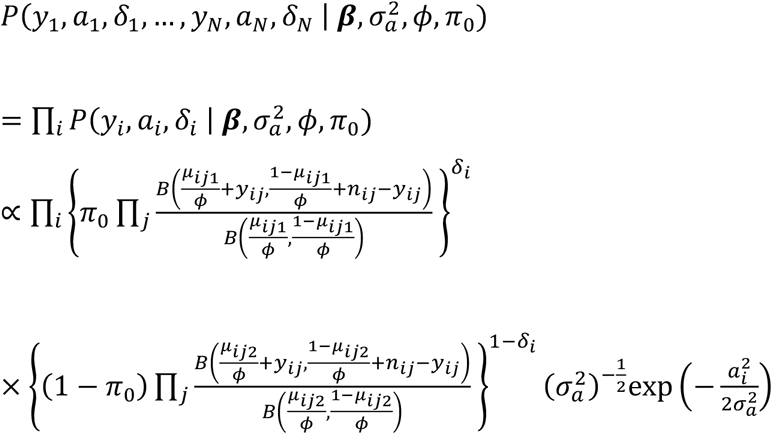

Here 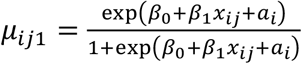 and 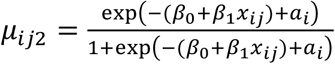. The variational EM iteration goes as follows:

In the E-step, we use variational inference^27,28^ to approximate the posterior distribution 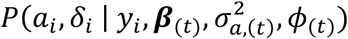, where 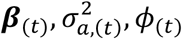 are the parameter values at the current iteration. We use the mean field approximation *q*(*a_i_, δ_i_*) = *q*(*a_i_*)*q*(*δ_i_*) with delta method approximation^27^. Denote the variational distribution by

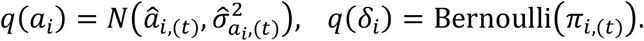

See **Supplementary Notes** for details of the derivation.

In the M-step, we first update *π*_0_ by 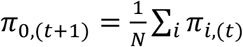 and update 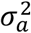 by 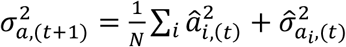. Update ***β*** and *ϕ* by numerical optimization of the following objective function:

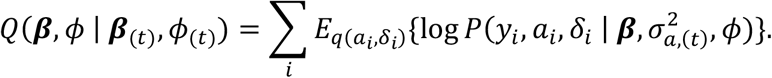

Here *E*_*q*(*a_i_,δ_i_*)_{·} is the expectation under variational distribution *q*(*a_i_, δ_i_*).

After the parameter estimation, we test the null hypothesis *H*_0_: *β*_1_ = 0 using likelihood ratio test. Rejecting this null hypothesis indicates that there is differential ASE with respect to the covariate.

### Simulation studies

We conduct simulation studies using total read counts and parameters estimated from a real endoderm differentiation dataset^10^. The dataset is comprised of 4,102 genes and 30,474 cells collected from 105 donors. See **Methods** subsection *Single-cell ASE data from endoderm differentiation* for details of the study. We randomly select 2,400 genes and use the real total allelespecific read counts as the total allele-specific read counts (*n_ij_*) in our simulations. This setting reflects realistic read depth and number of cells, but does not affect ASE which depends on the relative abundance of reference and alternative alleles. We simulate the alternative allele read counts assuming that there is only one eQTL driving the ASE

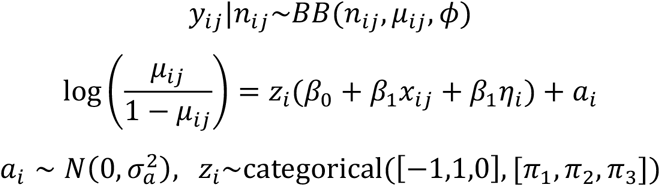

In contrast to the DAESC-Mix model, this simulation model introduces a third possible value of the latent variable *z_i_*. Besides two values −1 and 1 which are modeled by DAESC-Mix, the third value *z_i_* = 0 corresponds to the individuals for which the eQTL SNP is homozygous. The haplotype proportions *π*_1_, *π*_2_, *π*_3_ are simulated based on given LD coefficient (r^2^) between the eQTL and exonic SNP. We vary r^2^ to 0, 0.1 and 0.9, and simulate 800 genes for each value of r^2^ including 400 null genes and 400 non-null genes. The procedure to simulate the mixture probabilities with given r^2^ is described in the **Supplementary Notes**.

We include two covariates in the simulation to evaluate the performance of DAESC under two types of D-ASE. The continuous covariate *x_ij_* is the real pseudotime provided by the original study^10^; the discrete covariate *η_i_* is a simulated sample-level disease status which can take values 0 or 1. A randomly chosen half of the individuals are assigned *η_i_* = 0 (control) and the other half are assigned *η_i_* = 1 (case).

To choose realistic values of other parameters, we apply DAESC-BB to the real data and obtain estimates of *β*_0_, *β*_1_, 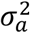 and *ϕ*. We select the genes with top 500 largest |*β*_1_| as potential values of parameters for the simulation. For each of the 2,400 genes, we randomly select a set of parameters 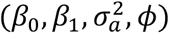 from the 500 candidate values. For null genes we reset *β*_1_ = 0. The 500 sets of candidate values are provided in **Supplementary Table 3** distribution of the parameters is visualized in **Supplementary Figure 6**.

We also vary the sample size to N=10, 50, 100. For D-ASE w.r.t. *x_ij_*, we randomly sample N individuals from the simulated data; for D-ASE w.r.t. *η_i_*, we randomly sample N/2 cases and N/2 controls.

### Simulations with multiple eQTL SNPs per gene

Due to the large number of scenarios for LD among eQTLs and the exonic SNP, we conduct this simulation study under a simplified scenario: all the eQTLs are independent from each other and independent from the exonic SNP. Similar to the one-eQTL scenario, we simulate the data using beta-binomial mixture model. Because the number of mixture components grow with the number of eQTLs, we simulate the mixture components indirectly, by simulating the genotypes of the eQTLs. The steps are as follows:

- Randomly choose 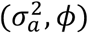 from 500 sets of candidate values (**Supplementary Table 3**). Parameters 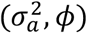 are the same across all mixture components.
- Simulate the minor allele frequency (MAF) of *m* eQTLs, from *MAF*_1_, *MAF*_2_, …, *MAF_m_*~Uniform[0.1,0.5].
- Simulate the alleles of eQTLs that resides on the haplotype of the <u>reference</u> allele of the exonic SNP for N individuals, denoted by *g*_*ik*0_~*bernoulli*(*MAF_k_*), *i* = 1, …, *N*; *k* = 1, …, *m*.
- Simulate the alleles of eQTLs that resides on the haplotype of the <u>alternative</u> allele of the exonic SNP, denoted by *g*_*ik*1_, *i* = 1, …, *N*; *k* = 1, …, *m*.
- Draw *m* pairs of regression coefficients (*β*_0_, *β*_1_) from 500 sets of candidate values (**Supplementary Table 3**), denoted by (*β*_10_, *β*_11_), …, (*β*_*m*0_, *β*_*m*1_).
- Compute individual-specific ASE effects size as 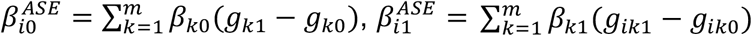.
- Compute *μ_ij_* from 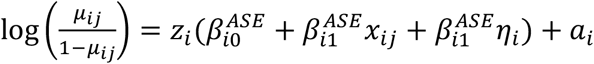. For individuals who have the same set of 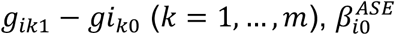 and 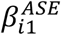 are the same and hence the model collapses into the beta-binomial mixture model.
- Generate *y_ij_~BB*(*n_ij_, μ_ij_, ϕ*).

We vary the number of eQTLs to *m* = 2, 3, 4, 5, 6.

### Other methods for comparison

We compare DAESC-BB and DAESC-Mix to two other methods. The first method is generalized linear mixed model (GLMM) implemented by the lme4 package in R. The GLMM is formulated as follows:

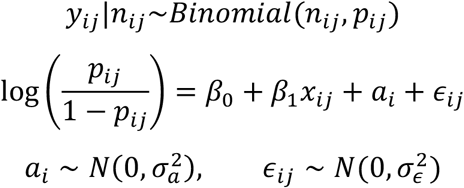

The R formula is cbind(y,n-y) ~ x + (1 |subj) + (1 |obs), where subj is the individual ID and obs is the unique ID for each cell. Here *α_i_* accounts for sample repeat structure and *ϵ_ij_* accounts for overdispersion.

For differential ASE across disease status, we further compare with EAGLE, a method for bulk tissue ASE analysis assuming independence across samples. We aggregate cell from each individual into a pseudobulk sample by summing the alternative and total read counts. We then apply EAGLE to test for differential ASE using the pseudobulk samples.

### Single-cell ASE data from endoderm differentiation

Cuomo et al^10^ conducted an endoderm differentiation experiment of 125 induced pluripotent stem cell (iPSC) lines from the Human Induced Pluripotent Stem Cell initiative (HipSci). Gene expression was profiled at 4 differentiation times points using single-cell RNA-seq (Smart-seq2). We obtained SNP-level allele-specific read counts for 114 donors from (https://zenodo.org/record/3625024#.YnJ-ivPMKi4), and restrict to 105 individuals for which genotype data are available to us. We remove SNPs with low mappability (ENCODE 75-mer mappability < 1), and those with monoallelic expression to reduce the effect of potential genotyping error. Monoallelic expression is defined for each SNP in each individual by ALT/TOTAL<0.02 or ALT/TOTAL>0.98^18^, where ALT is the sum of alternative allele read counts for all cells from the individual, and TOTAL is the corresponding sum of total allelespecific read counts.

### Aggregating SNP-level ASE counts to gene-level

Since phased genotype data are needed to aggregate SNP-level ASE counts to gene-level ASE counts, we impute and phase the genotype data using the Michigan Imputation Server with the Haplotype Reference Consortium (HRC) r1.1 data as the reference panel. For each individual and each gene, we sum the ASE counts across all SNPs within the exonic regions of the gene for each haplotype and obtain two haplotype-specific counts (“hap1” count and “hap2” count). The exonic regions are provided by GTEx v7^29^ annotation files (hg19) based on collapsed gene model. After removing the genes which have non-zero ASE counts in ≤ 20% of the cells, we obtain ASE counts for 4,102 genes and 30,474 cells.

For joint analysis across individuals, “alternative” and “reference” haplotypes need to be consistently assigned across individuals. In the paper by Cuomo et al^10^, the haplotype which is on the same chromosome as the alternative allele of the eQTL is assigned as the alternative haplotype. However, we would like to conduct ASE analysis without calling eQTL first, as is the case in many other studies. Therefore, we assign alternative and reference haplotypes based on the exonic SNP which has the highest total allele-specific read count across individuals (referred to by “top exonic SNP”), i.e. the haplotype on the same chromosome as the alternative allele of the top exonic SNP is assigned as the alternative haplotype. For those individuals for which the top exonic SNP is homozygous, alternative and reference haplotypes are assigned randomly.

### Comparing DAESC-Mix mixture labels and observed haplotype combinations

Since phased genotype data are available for this study, we can use them to validate the ability of DAESC-Mix to capture haplotype combinations. For each gene, we obtain a posterior probability (p_mix_) for each individual to belong to the first group. We assign the individual to the first group if p_mix_>0.5, or the second group if p_mix_<0.5. To compare with observed haplotype combinations, we first identify the top eQTL reported by Cuomo et al for each of the genes above. The original paper identified eQTL for three cell types separately: iPSC, mesendoderms and definitive endoderms. We choose the SNP that shows the strongest association p-value in any of the three cell types as the top eQTL for the gene. There are three possible observed haplotype combinations: 1) alt_eQTL_,alt_gene_|ref_eQTL_,ref_gene_, 2) alt_eQTL_,ref_gene_|ref_eQTL_,alt_gene_, 1) alt_eQTL_,alt_gene_|alt_eQTL_,ref_gene_ or ref_eQTL_,alt_gene_|ref_eQTL_,ref_gene_. Here ref_eQTL_ and alt_eQTL_ are the reference and alternative alleles of the top eQTL, respectively; ref_gene_ and alt_gene_ are the reference and alternative haplotypes of the gene, respectively. Alleles or haplotypes on same side of “|” are on the same haplotype. We tally the number of individuals in two mixture groups vs. three haplotype combinations into a 2×3 table (**Figure 3**). Finally, we perform Fisher’s exact test on the 2×3 table to test the association between mixture clusters and observed haplotype combinations.

### Dynamic eGene Clustering

We explore the total expression trends of (1) previously discovered dynamic eGenes by Cuomo et al^10^ and (2) the set of dynamic ASE genes discovered using DAESC-Mix (**Supplementary Table 1**). Pseudotime smoothing was performed as in Cuomo et al^10^, and spectral clustering was performed on pseudotime-smoothed total expression using Pearson correlation as the affinity metric. In order to maintain a meaningful comparison with the original analysis, 4 clusters were used for both analyses.

### Chromatin state analysis

We download the chromatin states learned by ChromHMM^20^ for the Roadmap Epigenomics Project^21^ (https://egg2.wustl.edu/roadmap/web_portal/chr_state_learning.html). For each gene, we compare the chromatin state at the TSS between iPSCs and endoderms. We consider chromatin states ≤ 7 as active, including 1_TssA, 2_TssAFlnk, 3_TxFlnk, 4_Tx, 5_TxWk, 6_EnhG, and 7_Enh, and assign them value 1 to represent active states in general. The remaining states are considered inactive and assigned value 0. Since there are multiple epigenomics for iPSCs (E018-E022, https://docs.google.com/spreadsheets/d/1yikGx4MsO9Ei36b64yOy9Vb6oPC5IBGlFbYEt-N6gOM/edit#gid=15), we use the average chromatin states (0 to 1) as the chromatin state for iPSC. We then compute the absolute difference of chromatin state between iPSC vs. hESC derived CD184+ endoderm cultured cells (E011), which we refer to as chromatin state change.

For three D-ASE methods, DAESC-BB, DAESC-Mix and GLMM, we compute the average chromatin state change for D-ASE genes (FDR<0.05) and non-D-ASE genes (FDR≥0.05), respectively. There are 324 D-ASE genes and 3,778 non-D-ASE genes identified by DAESC-BB, 657 D-ASE genes and 3,445 non-D-ASE genes identified by DAESC-Mix, and 1,995 D-ASE genes and 2,107 non-D-ASE genes identified by GLMM. To test the significance of the difference between D-ASE and non-D-ASE genes, we use linear regression adjusting for the total number of allele-specific reads for each gene: chromatin state change ~ I(D-ASE) + total read depth of the gene. This adjustment removes the effect of total expression, which can be a potential confounder. We also compute the correlation between D-ASE effect size (*β*_1_) and chromatin state change.

### Gene-set enrichment

We conduct gene set enrichment analysis for 657 D-ASE genes identified by DAESC-Mix using FUMA GWAS^30^. We only consider Gene Ontology (GO) biological process pathways^31^ and use protein-coding genes as background. Finally, gene sets with enrichment adjusted p-value <0.05 are considered as significantly enriched.

### Classification of dynamic ASE genes

We classify the D-ASE genes identified by DAESC-Mix based on the stage of differentiation where allelic imbalance occurs. For each D-ASE gene, we first compute the average allelic fraction for iPSCs (*p_ipsc_*) and definitive endoderms (*p_defendo_*) estimated by DAESC-Mix as 1/(1 + exp(−(*β*_0_ + *β*_1_*t*))), where *t* is the average pseudotime of the cell type. See Cuomo et al^10^ for the classification of cell types. Genes are classified into the following categories based on their ASE patterns:

- Increasing: *p_defendo_* < *p_ipsc_* < 0.47 or *p_defendo_* > *p_ipsc_* > 0.53.
- Decreasing: *p_ipsc_* < *p_defendo_* < 0.47 or *p_ipsc_* > *p_defendo_* > 0.53.
- Late: *|p_ipsc_* – 0.5| < 0.03 and *|p_defendo_* – 0.5| > 0.03
- Early: *|p_ipsc_* – 0.5| > 0.03 and *|p_defendo_* – 0.5| < 0.03
- Switching: *p_ipsc_* < 0.47 and *p_defendo_* > 0.53, or *p_defendo_* < 0.47 and *p_ipsc_* > 0.53

Other genes are classified as unspecified.

### Pancreatic islet data

Segerstolpe et al^8^ collected scRNA-seq data from pancreatic islet samples of 4 type 2 diabetes (T2D) patients and 6 controls. Libraries were prepared using Smart-seq2 protocols and sequencing was conducted using single-end 43 bp reads. We downloaded raw fastq files from ArrayExpress and trimmed the reads with trimmomatic v0.38^32^. We then aligned the reads to hg19 reference genome using STAR 2.7.10a^33^. We then marked duplicated reads with Picard 2.18.

Before obtaining ASE counts call, we first call genetic variants from scRNA-seq data using GATK (4.0.0). We followed the GATK best practices workflow for RNAseq short variant discovery. After further preprocessing steps (SplitNCigarReads and base recalibration), we merge the bam files of all cells from each individual into a pseudobulk bam file per individual. We then call variants using GATK HaplotyperCaller with the 10 pseudobulk bam files as input. We extract biallelic SNPs from the called variants. We then obtain single-cell ASE counts using GATK ASEReadCounter. We only retain the 2,209 cells that passed quality in the original paper^8^ and discard the rest.

For each individual, we remove SNPs with potential genotyping error. Specifically, we remove SNPs with genotyping read depth ≤ 10 and genotyping quality ≤ 15. We further remove the SNPs with monoallelic expression, defined by pseudobulk allelic fraction <0.05 or >0.95. The pseudobulk allelic fraction is defined as 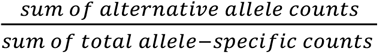, where the sums are taken across cells from the individual. This step is to further remove genotyping error.

To reduce the effect of alignment errors, we remove the SNPs with ENCODE 40-mer mappability <1. We then aggregate ASE counts from SNP level to gene level using a pseudo phasing approach used by the ASEP paper^14^. This pseudo phasing approach was performed on four major endocrine cells: alpha, beta, gamma and delta cells. We aggregate ASE counts from these four cell types into pseudobulk ASE counts. If there are multiple heterozygous exonic SNP within a gene, we sum the counts for the expression minor allele (the one with lower allele-specific read count) of all exonic SNPs as the alternative haplotype read count for the gene.

For cell-type-specific D-ASE analysis, we only analyzed genes that are available for a reasonably large number of cells and individuals. For each gene, we first remove individuals with <3 cells or <5 reads from the cell type. We drop the gene from D-ASE analysis if there are <50 cells or <2 cases or <2 controls remaining.

## Data and code availability

The DAESC R package and other analysis scripts is available on github: https://github.com/gqi/DAESC. The ASE data from endoderm differentiation is available on https://zenodo.org/record/3625024#.YnJ-ivPMKi4. Other HipSci data are available on https://www.hipsci.org/. The pancreatic islet data are available on ArrayExpress via accession number E-MTAB-5061.

## URLs

HipSci: https://www.hipsci.org/

ArrayExpress: https://www.ebi.ac.uk/arrayexpress/

ENCODE mappability: https://genome.ucsc.edu/cgi-bin/hgFileUi?db=hg19&g=wgEncodeMapability

Trimmomatic: http://www.usadellab.org/cms/?page=trimmomatic

STAR: https://github.com/alexdobin/STAR

Picard: https://broadinstitute.github.io/picard/

GATK: https://gatk.broadinstitute.org/hc/en-us

GATK Best Practices Workflows: https://gatk.broadinstitute.org/hc/en-us/sections/360007226651-Best-Practices-Workflows

