## Supplementary Figures for "Single-cell allele-specific expression analysis reveals dynamic and cell-type-specific regulatory effects"

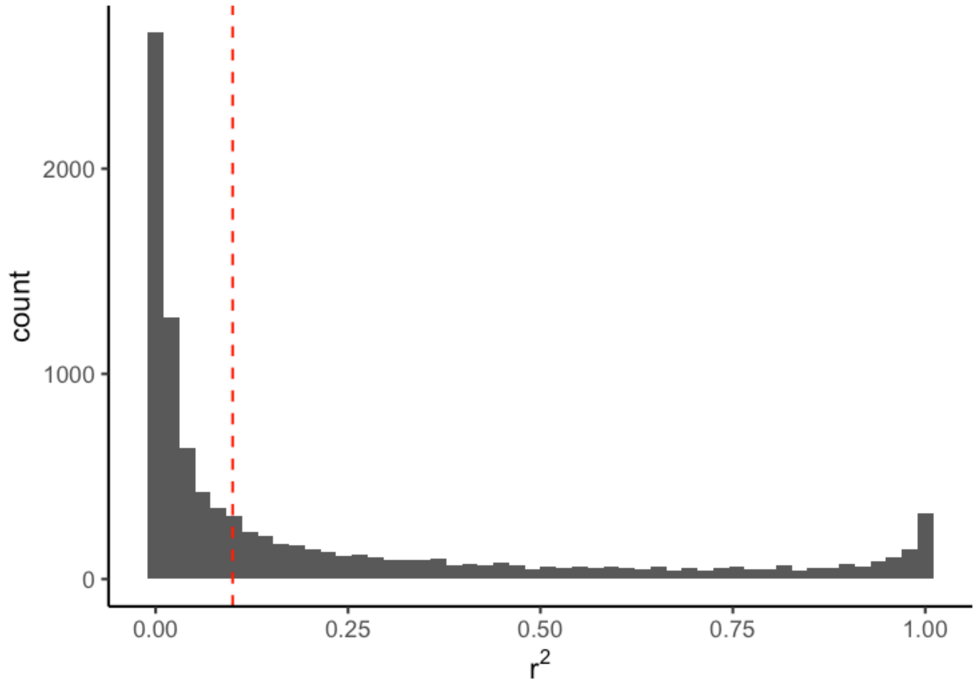


**Supplementary Figure 1. Distribution of linkage disequilibrium (LD) coefficient between the top eQTL and top exonic SNP of an eGene observed in GTEx v8 whole blood.** The top exonic SNP is defined as the exonic SNP with highest allele-specific read depth. The red dashed line is $r^2$=0.1 and 56.7% eQTL-exonic SNP pairs have $r^2$<0.1.

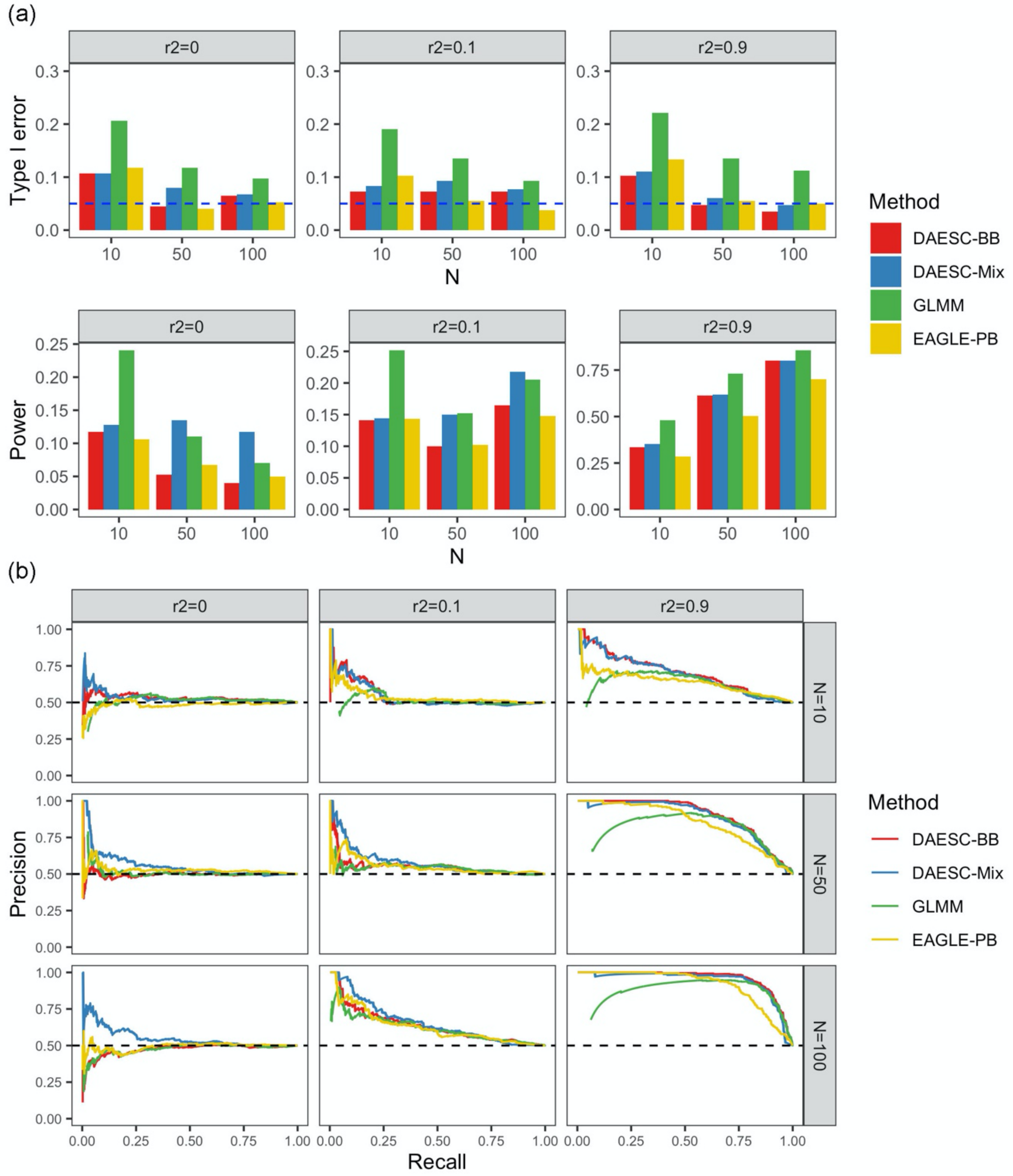


**Supplementary Figure 2. Performance of four methods for differential ASE detection of case vs control observed in simulation studies where one eQTL drives ASE.** (a) Type I error and power under significance threshold p<0.05 and (b) precision-recall curves. Allele-specific read counts are simulated from beta-binomial mixture model under the scenario where only one eQTL drives the ASE of an exonic SNP (see Methods). The linkage disequilibrium between the eQTL and the exonic SNP is varied to $r^2 = 0, 0.1, 0.9$, and the sample size is varied N=10, 50, 100. The number of cases and controls are both N/2.

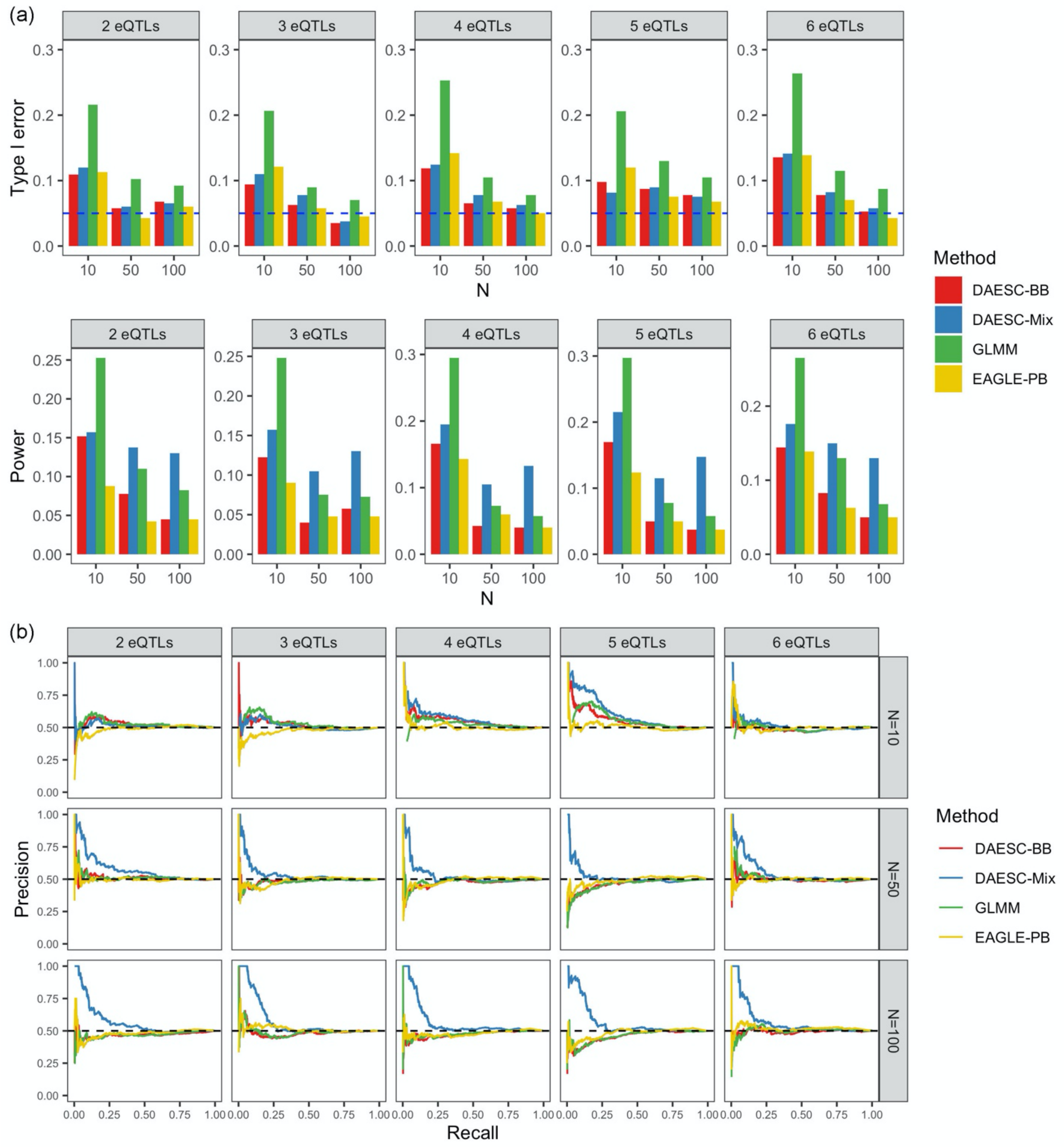


**Supplementary Figure 3. Performance of four methods for differential ASE detection of case vs control observed in simulation studies where multiple eQTLs drive ASE.** (a) Type I error and power under significance threshold $p<0.05$ and (b) precision-recall curves. Allele-specific read counts are simulated from beta-binomial mixture model assuming multiple eQTLs drive the ASE of an exonic SNP. The sample size is varied to N=10, 50, 100. The number of cases and controls are both N/2.

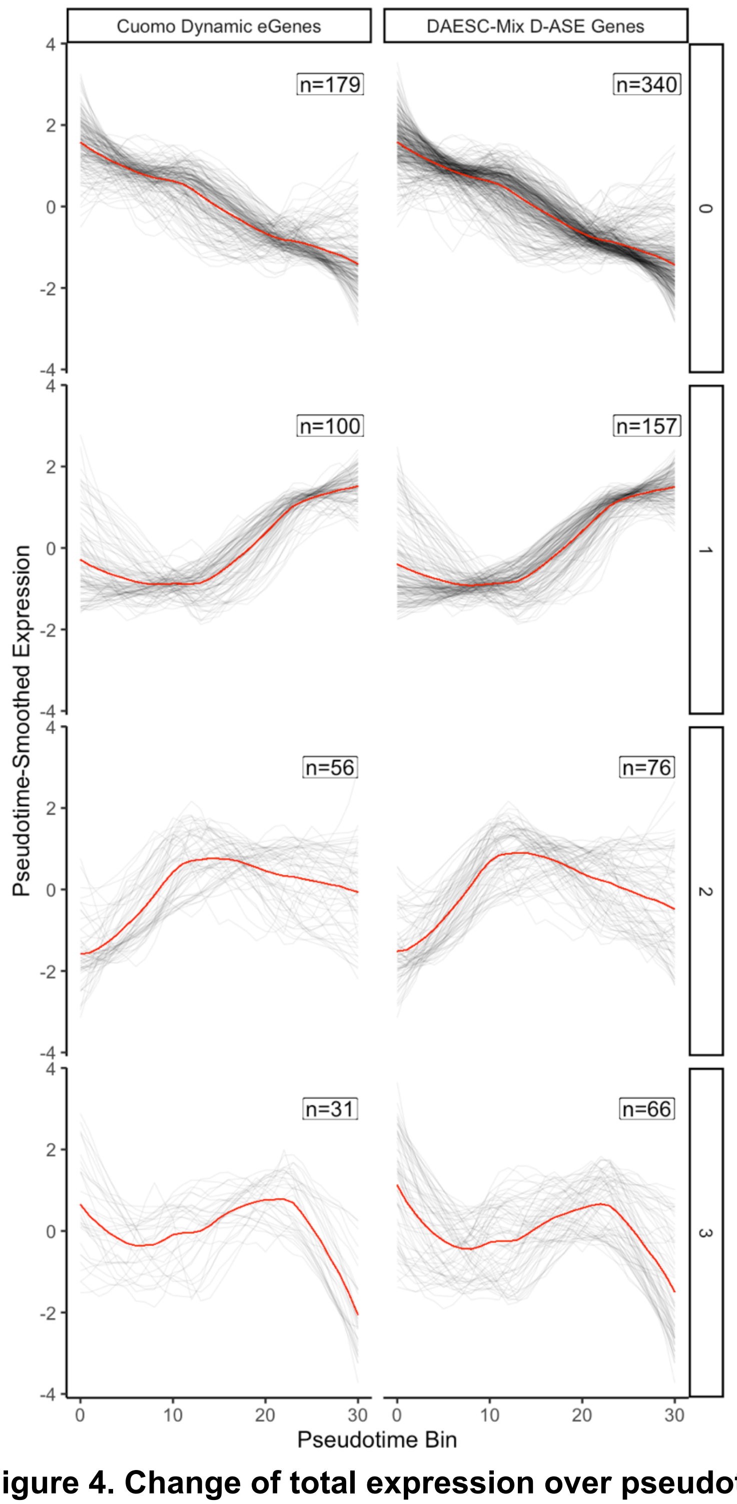


**Supplementary Figure 4. Change of total expression over pseudotime for D-ASE genes identified by DAESC-Mix and dynamic eGenes reported by Cuomo et al.** Each row is a cluster of patterns identified by spectral clustering. Each grey line is the trajectory of one gene and red line is the average trajectory within the cluster.

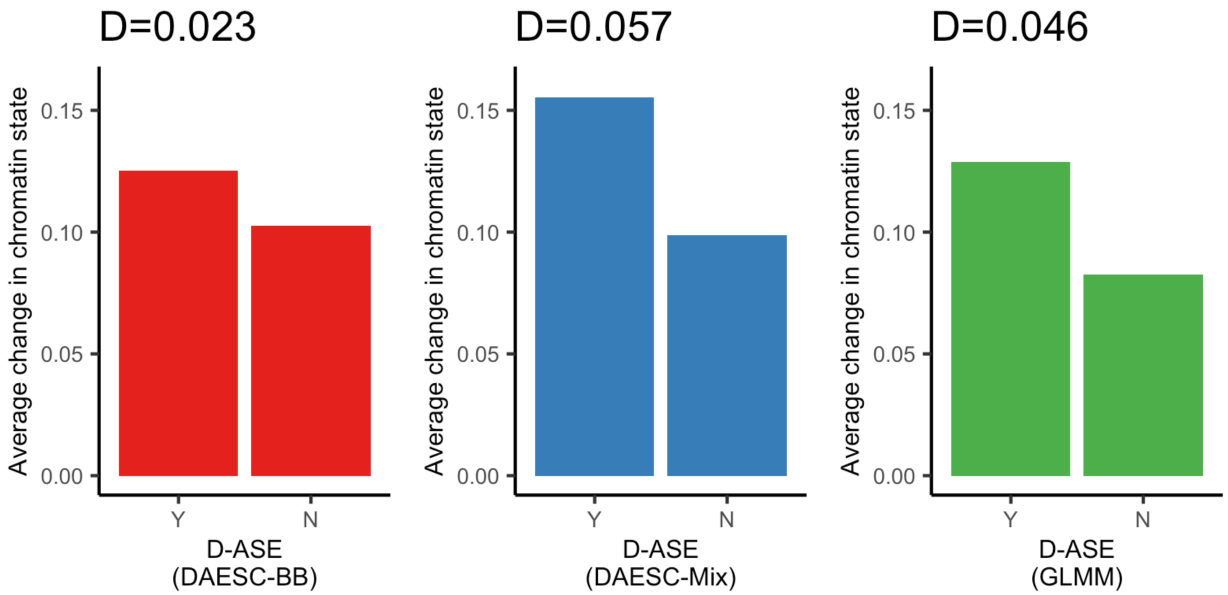


**Supplementary Figure 5. Average change of chromatin state of top 300 D-ASE genes (lowest p-value) during endoderm differentiation identified by three methods.** For each method, the top 300 genes (Y) are compared with 300 background genes (N) with matched total read depth. The difference between Y and N groups (D) for each method are shown above the panel.

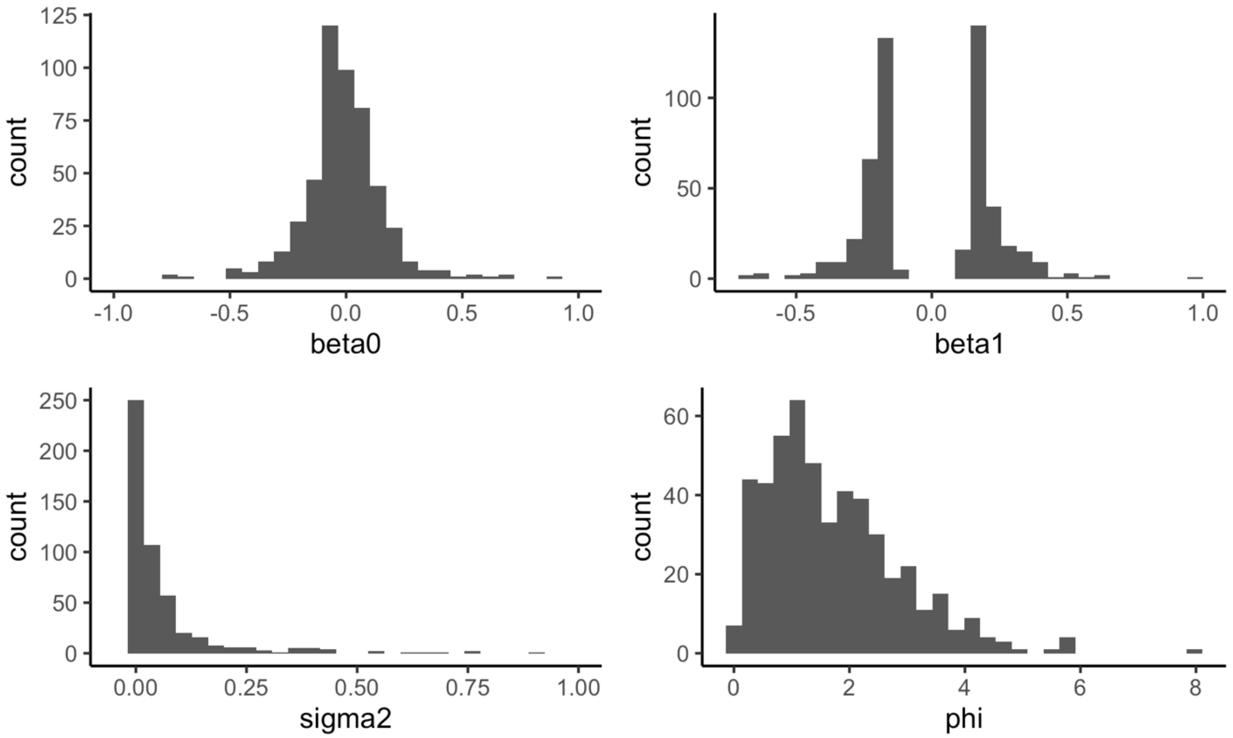


**Supplementary Figure 6. Distribution of simulation parameters.** Exact parameters are provided in Supplementary Table 3.
