## Supplementary Notes for "Single-cell allele-specific expression analysis reveals dynamic and cell-type-specific regulatory effects"

#### 1 DAESC-BB model and inference

##### 1.1 Model setup

For a gene or heterozygous exonic SNP, let $y_{ij}$ be the alternative allele read count of individual $i$ and cell $j$, and $n_{ij}$ be the total allele-specific read count. Here $i = 1, 2, ..., N$ and $j = 1, 2, ..., J_i$. Let $\boldsymbol{x}_{ij}$ be a length-$p$ vector of independent variables. The DAESC-BB model is formulated as follows:

$$y_{ij}|n_{ij} \sim \text{beta-binomial}(n_{ij}, \mu_{ij}, \phi) \tag{1}$$

$$\log(\mu_{ij}/(1-\mu_{ij})) = \boldsymbol{x}_{ij}^T\boldsymbol{\beta} + a_i \tag{2}$$

$$a_i \sim N(0, \sigma_a^2) \tag{3}$$

The fixed effects $\boldsymbol{\beta}$ represents ASE and dynamic ASE effects. The individual-specific random effects $a_i$ capture the sample repeat structure due to having multiple cells per individual.

##### 1.2 Parameter estimation by variational EM algorithm

We employ the variational EM algorithm (Wang and Blei, 2013; Blei et al, 2017) to estimate unknown parameters in model (1)-(2), treating $a_i$ as missing data. Define $\boldsymbol{y}_i = (y_{i1}, ..., y_{iJ_i})^T$ and $\sigma(x) = 1/(1 + \exp(-x))$ as the sigmoid function.

The complete data likelihood is

$$\begin{aligned}
&P(\boldsymbol{y}_1, a_1, ..., \boldsymbol{y}_N, a_N \mid \boldsymbol{\beta}, \sigma_a^2, \phi)\\
=&\prod_i \Big[\prod_j \binom{n_{ij}}{y_{ij}} \frac{B\big(\sigma(\boldsymbol{x}_{ij}^T\boldsymbol{\beta}+a_i)/\phi + y_{ij}, \sigma(\boldsymbol{x}_{ij}^T\boldsymbol{\beta}+a_i)/\phi + n_{ij} - y_{ij}\big)}{B\big(\sigma(\boldsymbol{x}_{ij}^T\boldsymbol{\beta}+a_i)/\phi, \sigma(\boldsymbol{x}_{ij}^T\boldsymbol{\beta}+a_i)/\phi\big)}\Big](2\pi\sigma_a^2)^{-\frac{1}{2}}\exp(-\frac{a_i^2}{2\sigma_a^2})\\
\propto&(\sigma_a^2)^{-\frac{N}{2}}\prod_i \Big[\prod_j \frac{B\big(\sigma(\boldsymbol{x}_{ij}^T\boldsymbol{\beta}+a_i)/\phi + y_{ij}, \sigma(\boldsymbol{x}_{ij}^T\boldsymbol{\beta}+a_i)/\phi + n_{ij} - y_{ij}\big)}{B\big(\sigma(\boldsymbol{x}_{ij}^T\boldsymbol{\beta}+a_i)/\phi, \sigma(\boldsymbol{x}_{ij}^T\boldsymbol{\beta}+a_i)/\phi\big)}\Big]\exp(-\frac{a_i^2}{2\sigma_a^2})
\end{aligned}$$

Hence the complete-data log-likelihood is

$$\begin{aligned}
&\log P(\boldsymbol{y}_1, a_1, ..., \boldsymbol{y}_N, a_N \mid \boldsymbol{\beta}, \sigma_a^2, \phi)\\
=&const - \frac{N}{2}\log(\sigma_a^2) - \frac{1}{2\sigma_a^2}\sum_i a_i^2\\
+&\sum_i\sum_j \log B\big(\sigma(\boldsymbol{x}_{ij}^T\boldsymbol{\beta}+a_i)/\phi + y_{ij}, \sigma(\boldsymbol{x}_{ij}^T\boldsymbol{\beta}+a_i)/\phi + n_{ij} - y_{ij}\big) - \log B\big(\sigma(\boldsymbol{x}_{ij}^T\boldsymbol{\beta}+a_i)/\phi, \sigma(\boldsymbol{x}_{ij}^T\boldsymbol{\beta}+a_i)/\phi\big)
\end{aligned}$$

##### 1.2.1 E-step

At iteration $t$, we approximate the conditional distribution $P(a_i \mid \boldsymbol{y}_i; \boldsymbol{\beta}_{(t)}, \sigma^2_{a,(t)}, \phi_{(t)})$ by $N(\hat{a}_{i,(t)}, \hat{\sigma}^2_{a_i,(t)})$. See section 3 for derivation of the variational approximation.

Here $\boldsymbol{\beta}_{(t)}$, $\sigma^2_{a,(t)}$ and $\phi_{(t)}$ are the current values at iteration $t$. The Q-function can be computed as

$$
\begin{aligned}
&Q(\boldsymbol{\beta}, \sigma_a^2, \phi \mid \boldsymbol{\beta}_{(t)}, \sigma^2_{a,(t)}, \phi_{(t)}) \\
=&const - \frac{N}{2}\log(\sigma_a^2) - \frac{1}{2\sigma_a^2}\sum_i \hat{a}^2_{i,(t)} + \hat{\sigma}^2_{a_i,(t)} \\
+&\sum_m w_m \{ \sum_i \sum_j \log B\big(\sigma(\boldsymbol{x}_{ij}^T\boldsymbol{\beta} + \hat{a}_{i,(t)} + \hat{\sigma}_{a_i,(t)} z_m)/\phi + y_{ij}, \sigma(\boldsymbol{x}_{ij}^T\boldsymbol{\beta} + \hat{a}_{i,(t)} + \hat{\sigma}_{a_i,(t)} z_m)/\phi + n_{ij} - y_{ij}\big) \\
&- \log B\big(\sigma(\boldsymbol{x}_{ij}^T\boldsymbol{\beta} + \hat{a}_{i,(t)} + \hat{\sigma}_{a_i,(t)} z_m)/\phi, \sigma(\boldsymbol{x}_{ij}^T\boldsymbol{\beta} + \hat{a}_{i,(t)} + \hat{\sigma}_{a_i,(t)} z_m)/\phi\big) \}
\end{aligned}
$$

Here $(z_m, w_m)$, $m = 1, ..., M$ are the nodes and weights of a Gaussian-Hermite quadrature for standard normal distribution. In practice, we found $M = 3$ nodes is sufficient for approximating the Q function.

##### 1.2.2 M-step

The update for $\sigma_a^2$ has a closed form:

$$
\sigma^2_{a,(t+1)} = \frac{1}{N}\sum_i \hat{a}^2_{i,(t)} + \hat{\sigma}^2_{a_i,(t)}
$$

The update for $\boldsymbol{\beta}$ and $\phi$ is obtained by optimizing the Q function using Newton-Raphson.

### 2 DAESC-Mix model and inference

#### 2.1 Model setup

DAESC-Mix is an extension of DAESC-BB incorporating implicit haplotype phasing. The model is formulated as follows:

$$
\begin{aligned}
&y_{ij} | n_{ij} \sim \text{beta-binomial}(n_{ij}, \mu_{ij}, \phi) && (4)\\
&\log(\mu_{ij}/(1-\mu_{ij})) = (2\delta_i - 1)\boldsymbol{x}_{ij}^T\boldsymbol{\beta} + a_i && (5)\\
&a_i \sim N(0, \sigma_a^2), \quad \delta_i \sim \text{Bernoulli}(\pi_0) && (6)
\end{aligned}
$$

#### 2.2 Parameter estimation by variational EM algorithm

We treat $a_i$ and $\delta_i$ as missing data. The complete-data likelihood is

$$
\begin{aligned}
&P(\boldsymbol{y}_1, a_1, \delta_1, ..., \boldsymbol{y}_N, a_N, \delta_N \mid \boldsymbol{\beta}, \sigma_a^2, \phi, \pi_0)\\
=&\prod_i \Big[\pi_0 \prod_j \binom{n_{ij}}{y_{ij}} \frac{B\big(\sigma(\boldsymbol{x}_{ij}^T\boldsymbol{\beta}+a_i)/\phi + y_{ij}, \sigma(\boldsymbol{x}_{ij}^T\boldsymbol{\beta}+a_i)/\phi + n_{ij} - y_{ij}\big)}{B\big(\sigma(\boldsymbol{x}_{ij}^T\boldsymbol{\beta}+a_i)/\phi, \sigma(\boldsymbol{x}_{ij}^T\boldsymbol{\beta}+a_i)/\phi\big)}\Big]^{\delta_i} \times\\
&\Big[(1-\pi_0) \prod_j \binom{n_{ij}}{y_{ij}} \frac{B\big(\sigma(-\boldsymbol{x}_{ij}^T\boldsymbol{\beta}+a_i)/\phi + y_{ij}, \sigma(-\boldsymbol{x}_{ij}^T\boldsymbol{\beta}+a_i)/\phi + n_{ij} - y_{ij}\big)}{B\big(\sigma(-\boldsymbol{x}_{ij}^T\boldsymbol{\beta}+a_i)/\phi, \sigma(-\boldsymbol{x}_{ij}^T\boldsymbol{\beta}+a_i)/\phi\big)}\Big]^{1-\delta_i} (2\pi\sigma_a^2)^{-\frac{1}{2}} \exp(-\frac{a_i^2}{2\sigma_a^2})\\
\propto&(\sigma_a^2)^{-\frac{N}{2}} \prod_i \Big[\pi_0 \prod_j \frac{B\big(\sigma(\boldsymbol{x}_{ij}^T\boldsymbol{\beta}+a_i)/\phi + y_{ij}, \sigma(\boldsymbol{x}_{ij}^T\boldsymbol{\beta}+a_i)/\phi + n_{ij} - y_{ij}\big)}{B\big(\sigma(\boldsymbol{x}_{ij}^T\boldsymbol{\beta}+a_i)/\phi, \sigma(\boldsymbol{x}_{ij}^T\boldsymbol{\beta}+a_i)/\phi\big)}\Big]^{\delta_i} \times\\
&\Big[(1-\pi_0) \prod_j \frac{B\big(\sigma(-\boldsymbol{x}_{ij}^T\boldsymbol{\beta}+a_i)/\phi + y_{ij}, \sigma(-\boldsymbol{x}_{ij}^T\boldsymbol{\beta}+a_i)/\phi + n_{ij} - y_{ij}\big)}{B\big(\sigma(-\boldsymbol{x}_{ij}^T\boldsymbol{\beta}+a_i)/\phi, \sigma(-\boldsymbol{x}_{ij}^T\boldsymbol{\beta}+a_i)/\phi\big)}\Big]^{1-\delta_i} \exp(-\frac{a_i^2}{2\sigma_a^2})
\end{aligned}
$$

The complete-data log-likelihood is

$$
\begin{aligned}
&\log P(\boldsymbol{y}_1, a_1, \delta_1, ..., \boldsymbol{y}_N, a_N, \delta_N \mid \boldsymbol{\beta}, \sigma_a^2, \phi, \pi_0)\\
=&const - \frac{N}{2}\log(\sigma_a^2) + \sum_i \Big\{ - \frac{a_i^2}{2\sigma_a^2}\\
&+\delta_i \Big[\log \pi_0 + \sum_j \log B\big(\sigma(\boldsymbol{x}_{ij}^T\boldsymbol{\beta}+a_i)/\phi + y_{ij}, \sigma(\boldsymbol{x}_{ij}^T\boldsymbol{\beta}+a_i)/\phi + n_{ij} - y_{ij}\big)\\
&- \sum_j \log B\big(\sigma(\boldsymbol{x}_{ij}^T\boldsymbol{\beta}+a_i)/\phi, \sigma(\boldsymbol{x}_{ij}^T\boldsymbol{\beta}+a_i)/\phi\big)\Big]\\
&+(1-\delta_i)\Big[\log(1-\pi_0) + \sum_j \log B\big(\sigma(-\boldsymbol{x}_{ij}^T\boldsymbol{\beta}+a_i)/\phi + y_{ij}, \sigma(-\boldsymbol{x}_{ij}^T\boldsymbol{\beta}+a_i)/\phi + n_{ij} - y_{ij}\big)\\
&- \sum_j \log B\big(\sigma(-\boldsymbol{x}_{ij}^T\boldsymbol{\beta}+a_i)/\phi, \sigma(-\boldsymbol{x}_{ij}^T\boldsymbol{\beta}+a_i)/\phi\big)\Big]\Big\}
\end{aligned}
$$

##### 2.2.1 E-step

At iteration $t$, we approximate the posterior distribution $P(a_i, \delta_i \mid \boldsymbol{y}_i; \boldsymbol{\beta}_{(t)}, \sigma^2_{a,(t)}, \phi_{(t)}, \pi_{0,(t)})$ using variational inference (Blei et al, 2017). Using the mean-field approximation $q(a_i, \delta_i) = q(a_i)q(\delta_i)$, we update $q(\delta_i)$ and $q(a_i)$ iteratively as follows. The update for $q(\delta_i)$ is

$$
\begin{aligned}
&\log q(\delta_i) = E_{q(a_i)}[\log P(a_i, \delta_i \mid \boldsymbol{y}_i, \boldsymbol{\beta}_{(t)}, \sigma^2_{a,(t)}, \phi_{(t)})]\\
=&const + \delta_i \Big[\log \pi_0 + \sum_j \int \log B\big(\sigma(\boldsymbol{x}_{ij}^T\boldsymbol{\beta}_{(t)}+a_i)/\phi_{(t)} + y_{ij}, \sigma(\boldsymbol{x}_{ij}^T\boldsymbol{\beta}_{(t)}+a_i)/\phi_{(t)} + n_{ij} - y_{ij}\big) q(a_i) da_i\\
&- \sum_j \int \log B\big(\sigma(\boldsymbol{x}_{ij}^T\boldsymbol{\beta}_{(t)}+a_i)/\phi_{(t)}, \sigma(\boldsymbol{x}_{ij}^T\boldsymbol{\beta}_{(t)}+a_i)/\phi_{(t)}\big) q(a_i) da_i\Big]\\
&+(1-\delta_i)\Big[\log(1-\pi_0) + \sum_j \int \log B\big(\sigma(-\boldsymbol{x}_{ij}^T\boldsymbol{\beta}_{(t)}+a_i)/\phi_{(t)} + y_{ij}, \sigma(-\boldsymbol{x}_{ij}^T\boldsymbol{\beta}_{(t)}+a_i)/\phi_{(t)} + n_{ij} - y_{ij}\big) q(a_i) da_i\\
&- \sum_j \int \log B\big(\sigma(-\boldsymbol{x}_{ij}^T\boldsymbol{\beta}_{(t)}+a_i)/\phi_{(t)}, \sigma(-\boldsymbol{x}_{ij}^T\boldsymbol{\beta}_{(t)}+a_i)/\phi_{(t)}\big) q(a_i) da_i\Big]
\end{aligned}
$$

The integrals are approximated by Gauss-Hermite quadrature. The resulting distribution is a bernoulli distribution, denoted by $\mathrm{Ber}(\pi_{i,(t)})$. The variational update for $a_i$ is

$$
\begin{aligned}
&\log q(a_i) = E_{q(\delta_i)}[\log P(a_i, \delta_i \mid \boldsymbol{y}_i, \boldsymbol{\beta}_{(t)}, \sigma^2_{a,(t)}, \phi_{(t)})] \\
=&const - \frac{a_i^2}{2\sigma^2_{a,(t)}} + \pi_{i,(t)}\Big[\sum_j \log B\big(\sigma(\boldsymbol{x}_{ij}^T\boldsymbol{\beta}_{(t)} + a_i)/\phi_{(t)} + y_{ij}, \sigma(\boldsymbol{x}_{ij}^T\boldsymbol{\beta}_{(t)} + a_i)/\phi_{(t)} + n_{ij} - y_{ij}\big) \\
-&\sum_j \log B\big(\sigma(\boldsymbol{x}_{ij}^T\boldsymbol{\beta}_{(t)} + a_i)/\phi_{(t)}, \sigma(\boldsymbol{x}_{ij}^T\boldsymbol{\beta}_{(t)} + a_i)/\phi_{(t)}\big)\Big] \\
+&(1 - \pi_{i,(t)})\Big[\sum_j \log B\big(\sigma(-\boldsymbol{x}_{ij}^T\boldsymbol{\beta}_{(t)} + a_i)/\phi_{(t)} + y_{ij}, \sigma(-\boldsymbol{x}_{ij}^T\boldsymbol{\beta}_{(t)} + a_i)/\phi_{(t)} + n_{ij} - y_{ij}\big) \\
-&\sum_j \log B\big(\sigma(-\boldsymbol{x}_{ij}^T\boldsymbol{\beta}_{(t)} + a_i)/\phi_{(t)}, \sigma(-\boldsymbol{x}_{ij}^T\boldsymbol{\beta}_{(t)} + a_i)/\phi_{(t)}\big)\Big]
\end{aligned}
$$

This update has no closed form, but we approximate by a normal distribution $N(\hat{a}_{i,(t)}, \hat{\sigma}^2_{a_i,(t)})$, as in Laplace variational inference (Wang and Blei, 2013). See section 3.2 for details.

##### 2.2.2 M-step

- Update $\pi_0$ by $\pi_{0,(t+1)} = \frac{1}{N}\sum\limits_i \pi_{i,(t)}$.
- Update $\sigma_a$ by $\sigma^2_{a,(t+1)} = \frac{1}{N}\sum_i \hat{a}^2_{i,(t)} + \hat{\sigma}^2_{a_i,(t)}$.
- Similar to section 1.2.2, update $\boldsymbol{\beta}$ and $\phi$ by numerical maximization of $E_{q(\delta_i,a_i)}[\log P(\boldsymbol{y}_1, a_1, \delta_1, ..., \boldsymbol{y}_N, a_N, \delta_N \mid \boldsymbol{\beta}, \sigma^2_a, \phi, \pi_0)]$, which is the complete data log likelihood integrated over variation distribution $q(\delta_i, a_i)$. The integration over $q(a_i)$ is conducted numerically using Gaussian-Hermite quadrature.

### 3 Approximating posterior distribution $P(a_i \mid \boldsymbol{y}_i)$ in the E-step

#### 3.1 DAESC-BB

In the DAESC-BB model, the joint distribution of $\boldsymbol{y}_i$ and $a_i$ is

$$
P(\boldsymbol{y}_i, a_i \mid \boldsymbol{\beta}, \sigma^2_a, \phi) \propto \prod_j \Big[\frac{B\big(\sigma(\boldsymbol{x}_{ij}^T\boldsymbol{\beta} + a_i)/\phi + y_{ij}, \sigma(\boldsymbol{x}_{ij}^T\boldsymbol{\beta} + a_i)/\phi + n_{ij} - y_{ij}\big)}{B\big(\sigma(\boldsymbol{x}_{ij}^T\boldsymbol{\beta} + a_i)/\phi, \sigma(\boldsymbol{x}_{ij}^T\boldsymbol{\beta} + a_i)/\phi\big)}\Big] \exp(-\frac{a_i^2}{2\sigma_a^2})
$$

Define $\boldsymbol{X}_i = (\boldsymbol{x}_{i1}, ..., \boldsymbol{x}_{iJ_i})^T$, $\boldsymbol{y}_i = (y_{i1}, ..., y_{iJ_i})^T$ and $\boldsymbol{n}_i = (n_{i1}, ..., n_{iJ_i})^T$. Define $f_{\boldsymbol{\beta},\sigma^2_a,\phi,\boldsymbol{X}_i,\boldsymbol{y}_i,\boldsymbol{n}_i}(a_i)$ as the log joint distribution, i.e.

$$
\begin{aligned}
f_{\boldsymbol{\beta},\sigma^2_a,\phi,\boldsymbol{X}_i,\boldsymbol{y}_i,\boldsymbol{n}_i}(a_i) =& \sum_j \log B\big(\sigma(\boldsymbol{x}_{ij}^T\boldsymbol{\beta} + a_i)/\phi + y_{ij}, [1 - \sigma(\boldsymbol{x}_{ij}^T\boldsymbol{\beta} + a_i)]/\phi + n_{ij} - y_{ij}\big) - \\
& \sum_j \log B\big(\sigma(\boldsymbol{x}_{ij}^T\boldsymbol{\beta} + a_i)/\phi, [1 - \sigma(\boldsymbol{x}_{ij}^T\boldsymbol{\beta} + a_i)]/\phi\big) - \frac{a_i^2}{2\sigma_a^2} \\
=& \sum_j \log B\big(\mu_{ij}/\phi + y_{ij}, (1 - \mu_{ij})/\phi + n_{ij} - y_{ij}\big) - \\
& \sum_j \log B\big(\mu_{ij}/\phi, (1 - \mu_{ij})/\phi\big) - \frac{a_i^2}{2\sigma_a^2} \\
=& \sum_j \log\Gamma\big(\mu_{ij}/\phi + y_{ij}\big) + \sum_j \log\Gamma\big((1 - \mu_{ij})/\phi + n_{ij} - y_{ij}\big) - \sum_j \log\Gamma\big(1/\phi + n_{ij}\big) - \\
& \sum_j \log\Gamma\big(\mu_{ij}/\phi\big) - \sum_j \log\Gamma\big((1 - \mu_{ij})/\phi\big) + J_i \log\Gamma\big(1/\phi\big) - \frac{a_i^2}{2\sigma_a^2}
\end{aligned}
$$

Here $\mu_{ij} = \sigma(\boldsymbol{x}_{ij}^T\boldsymbol{\beta} + a_i)$. To approximate $P(a_i \mid \boldsymbol{y}_i; \boldsymbol{\beta}, \sigma_a^2, \phi)$, we derived the Taylor expansion of $f_{\boldsymbol{\beta},\sigma_a^2,\phi,\boldsymbol{X}_i,\boldsymbol{y}_i,\boldsymbol{n}_i}(a_i)$. Denote by $\hat{a}_i$ the value that maximizes $f_{\boldsymbol{\beta},\sigma_a^2,\phi,\boldsymbol{X}_i,\boldsymbol{y}_i,\boldsymbol{n}_i}(a_i)$, we have

$$f_{\boldsymbol{\beta},\sigma_a^2,\phi,\boldsymbol{X}_i,\boldsymbol{y}_i,\boldsymbol{n}_i}(a_i) \approx f_{\boldsymbol{\beta},\sigma_a^2,\phi,\boldsymbol{X}_i,\boldsymbol{y}_i,\boldsymbol{n}_i}(\hat{a}_i) + \frac{1}{2} f''_{\boldsymbol{\beta},\sigma_a^2,\phi,\boldsymbol{X}_i,\boldsymbol{y}_i,\boldsymbol{n}_i}(\hat{a}_i)(a_i - \hat{a}_i)^2$$

Hence the posterior distribution $P(a_i \mid \boldsymbol{y}_i, \boldsymbol{\beta}, \sigma_a^2, \phi)$ can be approximated by $N(\hat{a}_i, \hat{\sigma}_{a_i}^2)$, where $\hat{\sigma}_{a_i}^2 = |f''_{\boldsymbol{\beta},\sigma_a^2,\phi,\boldsymbol{X}_i,\boldsymbol{y}_i,\boldsymbol{n}_i}(\hat{a}_i)|^{-1}$.

Now we derive the derivatives of $f_{\boldsymbol{\beta},\sigma_a^2,\phi,\boldsymbol{X}_i,\boldsymbol{y}_i,\boldsymbol{n}_i}(a_i)$ and the Newton-Raphson algorithm to obtain $\hat{a}_i$.

$$\begin{aligned} f'_{\boldsymbol{\beta},\sigma_a^2,\phi,\boldsymbol{X}_i,\boldsymbol{y}_i,\boldsymbol{n}_i}(a_i) =& \sum_j \Big[\psi\Big(\frac{\mu_{ij}}{\phi} + y_{ij}\Big) - \psi\Big(\frac{1-\mu_{ij}}{\phi} + n_{ij} - y_{ij}\Big) - \psi\Big(\frac{\mu_{ij}}{\phi}\Big) + \psi\Big(\frac{1-\mu_{ij}}{\phi}\Big)\Big]\mu_{ij}(1-\mu_{ij})/\phi - \frac{a_i}{\sigma_a^2} \\ f''_{\boldsymbol{\beta},\sigma_a^2,\phi,\boldsymbol{X}_i,\boldsymbol{y}_i,\boldsymbol{n}_i}(a_i) =& \sum_j \Big\{\Big[\psi_1\Big(\frac{\mu_{ij}}{\phi} + y_{ij}\Big) + \psi_1\Big(\frac{1-\mu_{ij}}{\phi} + n_{ij} - y_{ij}\Big) - \psi_1\Big(\frac{\mu_{ij}}{\phi}\Big) - \psi_1\Big(\frac{1-\mu_{ij}}{\phi}\Big)\Big]\mu_{ij}^2(1-\mu_{ij})^2/\phi^2 + \\ & \Big[\psi\Big(\frac{\mu_{ij}}{\phi} + y_{ij}\Big) - \psi\Big(\frac{1-\mu_{ij}}{\phi} + n_{ij} - y_{ij}\Big) - \psi\Big(\frac{\mu_{ij}}{\phi}\Big) + \psi\Big(\frac{1-\mu_{ij}}{\phi}\Big)\Big](1-2\mu_{ij})\mu_{ij}(1-\mu_{ij})/\phi\Big\} - \frac{1}{\sigma_a^2} \end{aligned}$$

Here $\psi(x) = \frac{d}{dx}\log\Gamma(x)$ is the digamma function and $\psi_1(x) = \frac{d^2}{dx^2}\log\Gamma(x)$ is the trigamma function. To obtain $\hat{a}_i$, we use a modified Newton-Raphson method. At iteration $k$, we update $a_i$ as follows

$$a_i^{k+1} = a_i^k - \tau f'(a_i^k)/f''(a_i^k)$$

By default, we set $\tau = 0.9$.

#### 3.2 DAESC-Mix

In the E-step for DAESC-Mix, the log variational distribution for $a_i$ is

$$\begin{aligned} h(a_i) =& \log q(a_i) \\ =& const - \frac{a_i^2}{2\sigma_a^2} + \pi_i\Big[\sum_j \log B\big(\sigma(\boldsymbol{x}_{ij}^T\boldsymbol{\beta} + a_i)/\phi + y_{ij}, \sigma(\boldsymbol{x}_{ij}^T\boldsymbol{\beta} + a_i)/\phi + n_{ij} - y_{ij}\big) \\ & - \sum_j \log B\big(\sigma(\boldsymbol{x}_{ij}^T\boldsymbol{\beta} + a_i)/\phi, \sigma(\boldsymbol{x}_{ij}^T\boldsymbol{\beta} + a_i)/\phi\big)\Big] \\ & + (1-\pi_i)\Big[\sum_j \log B\big(\sigma(-\boldsymbol{x}_{ij}^T\boldsymbol{\beta} + a_i)/\phi + y_{ij}, \sigma(-\boldsymbol{x}_{ij}^T\boldsymbol{\beta} + a_i)/\phi + n_{ij} - y_{ij}\big) \\ & - \sum_j \log B\big(\sigma(-\boldsymbol{x}_{ij}^T\boldsymbol{\beta} + a_i)/\phi, \sigma(-\boldsymbol{x}_{ij}^T\boldsymbol{\beta} + a_i)/\phi\big)\Big] \\ =& const + \pi_i f_{\boldsymbol{\beta},\sigma_a^2,\phi,\boldsymbol{X}_i,\boldsymbol{y}_i,\boldsymbol{n}_i}(a_i) + (1-\pi_i) f_{-\boldsymbol{\beta},\sigma_a^2,\phi,\boldsymbol{X}_i,\boldsymbol{y}_i,\boldsymbol{n}_i}(a_i) \end{aligned}$$

Here we dropped the subscript $(t)$ for simpler notations. The derivatives can be computed as follows:

$$\begin{aligned} h'(a_i) &= \pi_i f'_{\boldsymbol{\beta},\sigma_a^2,\phi,\boldsymbol{X}_i,\boldsymbol{y}_i,\boldsymbol{n}_i}(a_i) + (1-\pi_i) f'_{-\boldsymbol{\beta},\sigma_a^2,\phi,\boldsymbol{X}_i,\boldsymbol{y}_i,\boldsymbol{n}_i}(a_i) \\ h''(a_i) &= \pi_i f''_{\boldsymbol{\beta},\sigma_a^2,\phi,\boldsymbol{X}_i,\boldsymbol{y}_i,\boldsymbol{n}_i}(a_i) + (1-\pi_i) f''_{-\boldsymbol{\beta},\sigma_a^2,\phi,\boldsymbol{X}_i,\boldsymbol{y}_i,\boldsymbol{n}_i}(a_i) \end{aligned}$$

Similar to section 3.1, we derive the maximum $\hat{a}_i$ using Newton-Raphson updates

$$a_i^{k+1} = a_i^k - \tau h'(a_i^k)/h''(a_i^k)$$

and $q(a_i)$ can be approximated by $N(\hat{a}_i, \hat{\sigma}_{a_i}^2)$, where $\hat{\sigma}_{a_i}^2 = |h''(\hat{a}_i)|^{-1}$.

### 4 Simulate haplotype proportions

In the simulation studies, we vary the LD coefficient ($r^2$) between the eQTL and the exonic SNP (eSNP). The simulation model, however, does not directly use the LD coefficient. Instead, it uses haplotype proportions determined by $r^2$ and the minor allele frequencies (MAF).

We start by introducing a few notations. Denote by $g_{r1}$ the genotype of eQTL (regulatory SNP) in haplotype 1 and $g_{r2}$ the genotype of eQTL in haplotype 2. Similarly define $g_{e1}$ and $g_{e2}$ as the genotypes of eSNP in haplotypes 1 and 2, respectively. Genotypes $g_{r2}, g_{r2}, g_{e1}, g_{e2}$ can takes values 0 or 1. Denote by $a_r$ and $a_e$ the minor allele frequencies (MAF) of the eQTL and eSNP, respectively. We start by simulating $a_r$ from a uniform distribution: $a_r \sim U[0.1, 0.5]$.

With a given $r^2$, the possible values of $a_e$ are bounded by $a_r$. We derive the bound by first deriving the relationship among $r^2$, MAFs and haplotype frequencies. Note that

$$
\begin{aligned}
r =& \frac{E(g_{r1} + g_{r2} - Eg_{r1} - Eg_{r2})(g_{e1} + g_{e2} - Eg_{e1} - Eg_{e2})}{\sqrt{var(g_{r1} + g_{r2})var(g_{e1} + g_{e2})}} \\
=& \frac{E(g_{r1} + g_{r2} - 2a_r)(g_{e1} + g_{e2} - 2a_e)}{\sqrt{4a_r a_e(1 - a_r)(1 - a_e)}} \\
=& \frac{E(g_{r1}g_{e1}) + E(g_{r2}g_{e2}) - 2a_r a_e}{\sqrt{4a_r a_e(1 - a_r)(1 - a_e)}} \\
=& \frac{P(g_{r1} = 1, g_{e1} = 1) - a_r a_e}{\sqrt{a_r a_e(1 - a_r)(1 - a_e)}}
\end{aligned}
$$

Without loss of generality we assume $r > 0$. If $r < 0$ we can simply flip the reference and alternative alleles of one of the SNPs to ensure $r > 0$. Define the following haplotype frequencies:

$$
\begin{aligned}
p_{11} &= P(g_{r1} = 1, g_{e1} = 1), \quad p_{10} = P(g_{r1} = 1, g_{e1} = 0) \\
p_{01} &= P(g_{r1} = 0, g_{e1} = 1), \quad p_{00} = P(g_{r1} = 0, g_{e1} = 0)
\end{aligned}
$$

Hence $p_{11} = a_r a_e + r\sqrt{a_r a_e(1 - a_r)(1 - a_e)}$. It needs to satisfy the restrictions $p_{11} < a_r$ and $p_{11} < a_e$ since the haplotype frequency cannot exceed corresponding allele frequencies of individual SNPs. This is equivalent to

$$
r^2 \frac{a_e}{1 - a_e} \le \frac{a_r}{1 - a_r}, \quad r^2 \frac{a_r}{1 - a_r} \le \frac{a_e}{1 - a_e}.
$$

Hence we can derive the bounds for $a_e$:

$$
\frac{r^2 a_r}{1 - a_r + r^2 a_r} \le a_e \le \frac{a_r}{r^2(1 - a_r) + a_r}.
$$

We simulate $a_e$ by uniform distribution: $a_e \sim U[\frac{r^2 a_r}{1 - a_r + r^2 a_r}, \frac{a_r}{r^2(1 - a_r) + a_r}]$. Finally, we calculate the haplotype frequencies by

$$
\begin{aligned}
p_{11} =& a_r a_e + r\sqrt{a_r a_e(1 - a_r)(1 - a_e)} \\
p_{01} =& a_e - p_{11}, \; p_{10} = a_r - p_{11}, \; p_{00} = 1 - p_{11} - p_{01} - p_{10}.
\end{aligned}
$$

Hence the mixture probabilities are calculated by

$$\tilde{\pi}_1 = 2p_{00}p_{11},\ \tilde{\pi}_2 = 2p_{01}p_{10},\ \tilde{\pi}_3 = 2p_{10}p_{11} + 2p_{00}p_{01}.$$

These are the proportions of individuals for which the eQTL is heterozygous ($\tilde{\pi}_1, \tilde{\pi}_2$) or homozygous ($\tilde{\pi}_3$) in the general population, regardless of whether the eSNP is heterozygous. However, we need to restrict to the individuals for which the eSNP is heterozygous, since ASE cannot be measured for homozygous individuals. Hence we normalize the probabilities to get the final mixture probabilities:

$$\pi_k = \frac{\tilde{\pi}_k}{\tilde{\pi}_1 + \tilde{\pi}_2 + \tilde{\pi}_3}, \quad k = 1, 2, 3.$$
